# SpectroBird: Event-Level Vocalization Detection and Curation for Avian Ecology

**DOI:** 10.64898/2026.10.13.758621

**Authors:** Erik G. Dreyer, Pranav Gokhale, Vignesh Chandran, Tom Fox, Andrei Hushcha, Vanessa B. Prema, Charles R. Clark, Yasha Iravantchi, Marco Postiglione, Breanna Shi, Stephen Mussmann, Benjamin Freeman

## Abstract

Passive acoustic monitoring often summarizes long recordings as fixed-window presence predictions, but many ecological questions require records of individual vocal events: how many occurred, when they occurred, whether they overlapped, how long they lasted, what frequency band they occupied, and whether each event was complete enough for structural measurement. We address this event-level measurement problem with SpectroBird, a spectrogram-based method for localizing and curating avian vocalizations in noisy field recordings. SpectroBird first applies a detection-localization model to identify candidate vocalizations as time-frequency boxes. An event-consolidation step merges repeated detections from overlapping windows while preserving overlapping vocalizations as separate events. A completeness-curation model then classifies each consolidated event as complete or incomplete before downstream event-level analysis.

We demonstrate SpectroBird on breeding-season Hume’s Leaf Warbler buzz vocalizations recorded in the Himalaya across a 3,200–3,800 m elevational gradient. Under shared window-level scoring, the detection-localization model remained competitive with AST and YOLOv11-cls baselines while retaining event timing and frequency bounds; event-level evaluation achieved AP *>* 0.999 for complete buzzes. Applied to two field seasons, the resulting event records showed that duration of complete buzzes declined with elevation, complete-buzz occurrence showed no simple elevation trend, and relative day-of-year occurrence patterns were strongly correlated between years. These results show how event-level detection and curation can support ecological analyses that depend on individual vocalizations rather than fixed-window presence alone.

## 1. Introduction

Passive acoustic monitoring (PAM) is widely used for ecological observation, including biodiversity monitoring, phenology tracking, and habitat-use assessment, because it can collect long-duration audio across space and time with low disturbance (Sugai et al., 2019; Kahl et al., 2021). In practice, many PAM machine-learning pipelines return clip-level presence/absence or fixed-window presence predictions. These outputs are useful for occupancy and broad occurrence trends, but they do not directly measure individual vocal events. Many ecological questions instead ask how many vocalizations occurred, when they occurred, whether they overlapped, how long they lasted, and what frequency band they occupied. For such questions, fixed-window predictions must be converted into event-level records before they can support downstream ecological analysis.

To address this event-level measurement problem, we developed SpectroBird, a computer-vision method for spectrograms in which each candidate vocalization is treated as a localized object with explicit temporal and spectral extent. SpectroBird first applies a detection-localization model to identify candidate vocalizations as time-frequency boxes. An event-consolidation step merges repeated detections from overlapping windows while preserving overlapping vocalizations as separate events. A completeness-curation model then classifies each consolidated event as *complete* or *incomplete* before downstream event-level analysis. The detection-localization model is evaluated against benchmark full-window classifiers, including the Audio Spectrogram Transformer (AST) (Gong et al., 2021) and a YOLOv11-cls image-classification baseline (Khanam and Hussain, 2024).

Our main contributions are:

1. A practical method for short bioacoustic events that combines detection-localization, event consolidation, and completeness curation before down-stream ecological analysis.
2. A window-geometry-informed event-consolidation step that converts repeated detections from overlapping spectrogram windows into event-level records while helping preserve overlapping vocalizations as separate events.
3. A controlled comparison using a task-aligned spectrogram representation, showing that the detector remains competitive with full-window AST and YOLOv11-cls baselines under shared window-level scoring while retaining event timing and frequency bounds.
4. An end-to-end avian field application demonstrating that curated event records can support ecological analyses of occurrence, vocalization structure, and seasonal patterns.

We evaluate SpectroBird using buzz vocalizations of Hume’s Leaf Warbler (HLW; Fig. 1a; *Phylloscopus humei*), a high-elevation Himalayan passerine recorded near Sural village, Himachal Pradesh, India, across a 3,200–3,800 m elevational gradient (Fig. 2). Males produce a distinctive high-frequency buzz during the breeding season, likely in territorial and/or mate-attraction contexts (Fig. 1b). Our goal was to recover event-level records of these buzzes so occurrence, duration, and frequency placement could be summarized across recorder sites. In field recordings, these buzzes are embedded in variable background noise, and incompletely observed buzzes commonly arise under low SNR (e.g., distant birds) and transient non-stationary noise (Fig. 4). Buzzes can also overlap with other acoustic activity, including other HLW buzzes (Fig. 5). For this measurement goal, the method must localize buzzes in time-frequency space, preserve distinct overlapping events, and flag incompletely observed events before summary analysis. Otherwise, truncated or masked buzzes can underestimate event duration or frequency extent and bias downstream summaries.

**Figure 1:**
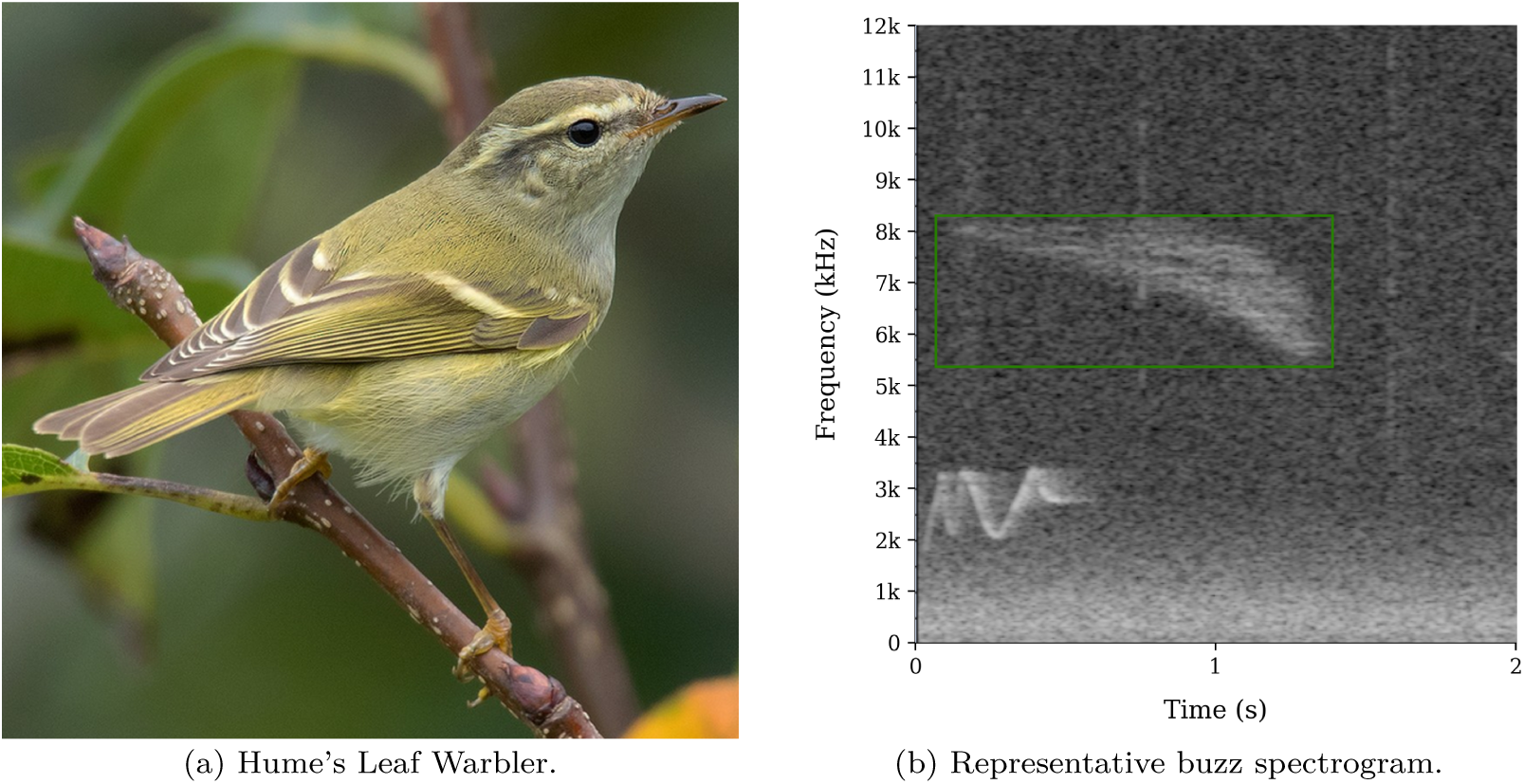
Study species and target vocalization. The Hume’s Leaf Warbler is a high-elevation Himalayan passerine; males produce a distinctive high-frequency buzz during the breeding season. The green box marks an annotated buzz used during initial data exploration.

**Figure 2:**
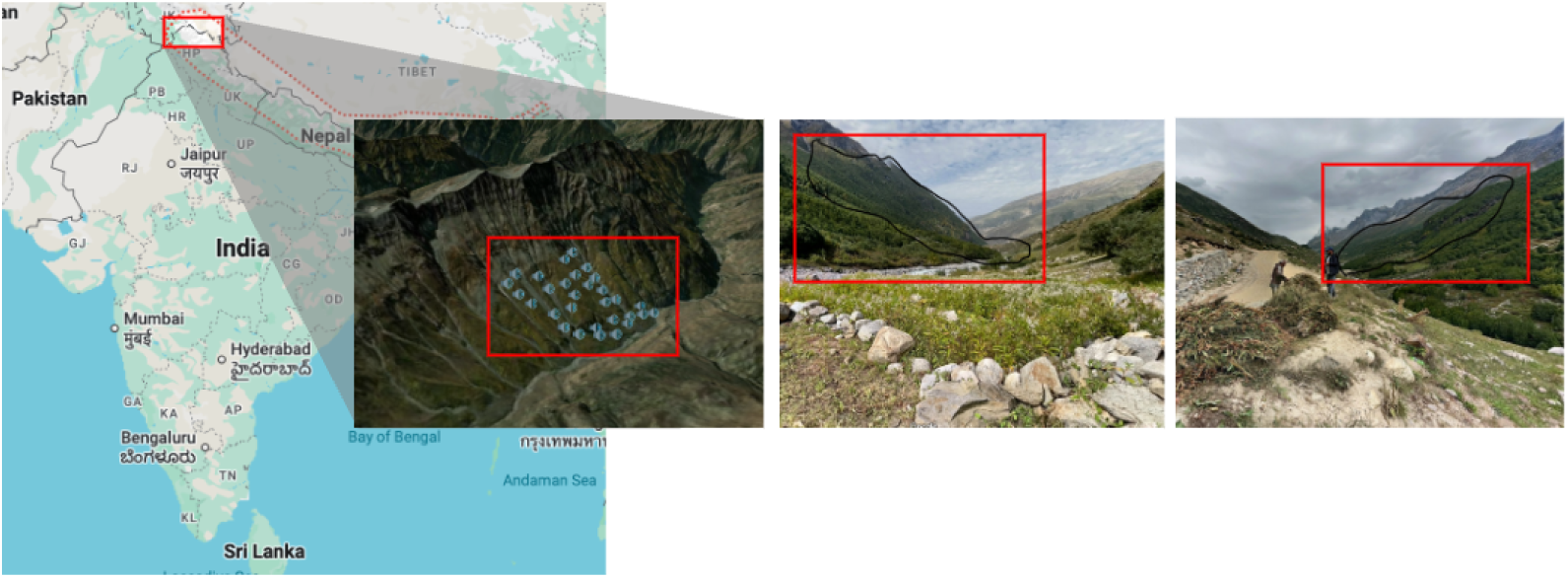
Study-area location in the Western Himalaya, India, an oblique topographic view of the recorder deployment area, and field photographs of recording sites, with the focal hillside outlined in red.

Using these event records, we analyzed occurrence and duration of retained complete buzzes across the elevational gradient. Environmental conditions across this gradient differ in ways that could affect breeding behavior, including warmer conditions, earlier breeding, and greater food availability at lower elevations than at colder, harsher upper elevations. We therefore hypothesized that lower elevations provide more favorable breeding conditions for HLW and derived two predictions. First, buzz vocalizations should occur more frequently at lower elevations, where interaction rates among rival males and/or between males and females may be higher. Second, buzz vocalizations should be longer at lower elevations, where males may be in better condition and therefore able to produce more exaggerated displays. We additionally compared complete-buzz occurrence between years and examined whether its day-of-year pattern was similar between years.

## 2. Related Work

### 2.1. Bioacoustic Monitoring and Species Detection

Passive acoustic monitoring (PAM) has become a core approach for biodiversity and behavioral studies because it enables repeated, low-disturbance sampling across large spatial and temporal scales (Sugai et al., 2019). In practice, however, operational PAM systems must handle limited labels, non-stationary background noise, overlap among vocal sources, and domain shift between training and deployment environments (Kahl et al., 2021). These constraints are especially relevant for species- and call-type-specific workflows, where target events are sparse and class boundaries can be subtle.

### 2.2. Clip-Level Classification for Bird Audio

Recent bird-audio pipelines have shown clip-level use of pretrained and fine-tuned models (including BirdNET and transformer-based approaches) (Kahl et al., 2021; Gong et al., 2021). These systems are typically used for presence/absence questions over fixed windows. For ecological tasks that depend on start time, end time, duration, or reliable frequency-band localization (e.g., call-bout structure analysis, or comparisons of frequency-band use with ecological correlates such as time-of-day, season, or elevation), additional event-level structure is needed because multiple events can occur within one window and event timing/frequency placement remain ambiguous. Many general audio classifiers use log-mel spectrogram representations designed for broad audio tagging, whereas the present task requires preserving fine frequency structure in a high-frequency band.

#### 2.2.1. BirdNET as a species-level reference model

BirdNET is widely used as a general-purpose multi-label bird-audio classifier in passive acoustic monitoring workflows (Kahl et al., 2021). We considered BirdNET as a contextual reference, but its outputs are not directly comparable to the buzz-specific benchmarks used here.

Three issues limit direct comparison. First, BirdNET is a species-level model: the HLW output node represents species-level acoustic evidence rather than the buzz call type specifically. HLW also produces non-buzz call vocalizations, which are common in the field recordings and were not annotated in our buzz-event datasets; a high BirdNET HLW score in a window without a labeled buzz could therefore reflect a true HLW call rather than a buzz false positive, making false-positive and true-negative counts undefined under the present labels. Second, BirdNET produces per-species confidence scores across many trained classes, and species-specific operating thresholds require separate calibration and validation before precision-recall values can be interpreted for a deployment setting (Schwinger et al., 2026). Third, Bird-NET uses a log-mel spectrogram representation, which allocates fewer bins per Hz at high frequencies than the linear high-band representation used here and may therefore reduce sensitivity to the frequency structure of HLW buzzes. Thus, BirdNET is best treated here as a species-level contextual model rather than a direct buzz-event benchmark; the quantitative benchmarks in this manuscript focus on models that can be trained and scored directly against buzz-specific window and event annotations.

### 2.3. Event-Level Detection and Localization

Event-level formulations address this boundary ambiguity by producing localized predictions over time and, in spectrogram object detection, over both time and frequency. Event-based matching and precision-recall analysis are formalized in the audio event detection literature (Mesaros et al., 2016) and are increasingly relevant to ecological audio-event detection workflows (Araya-Salas et al., 2023). Strongly annotated bird-audio corpora with species labels and start/end times provide individual vocalization intervals and have made time-localized event evaluation operationally feasible in bird-audio workflows (Gómez-Gómez et al., 2023). However, one-dimensional time labels do not provide frequency bounds and may be less suited to cases where concurrent or overlapping vocalizations must be separated into distinct event traces. Two vocalizations can share the same start and end times but occupy different frequency bands, so temporal labels alone cannot distinguish them as separate events. For HLW buzzes, this distinction matters because duration and frequency-band placement are both downstream measurements, and overlapping buzzes from different individuals can occur within the same recording window. The present task therefore uses two-dimensional time-frequency boxes to represent multiple candidate events with confidence and boundaries, supporting transparent post-processing and error analysis under overlap.

### 2.4. Windowed Object Detection and Box Consolidation

Object detection on overlapping image patches is a common computer-vision strategy when targets are small relative to the full image or when high-resolution inputs must be processed in manageable tiles. Slicing-aided inference frameworks such as SAHI improve small-object detection by running detectors on overlapping image patches and mapping predictions back to the original coordinate system (Akyon et al., 2022). The resulting duplicated or fragmented boxes are commonly handled with post-processing methods such as non-maximum suppression, Soft-NMS, and weighted box fusion (Bodla et al., 2017; Solovyev et al., 2021). More recent work on window-based object detection explicitly uses detection-window coordinates to suppress or merge redundant partial detections (Kos, 2025). Our consolidation step is related to this windowed-detection post-processing literature, but is adapted to acoustic spectrograms. Although overlap ambiguity is not unique to acoustic detection, it is especially consequential here. Overlapping boxes may reflect duplicate views of the same buzz across adjacent analysis windows, or distinct buzzes that genuinely overlap in time and frequency; overlap alone therefore cannot reliably determine whether detections should be merged. This motivates a consolidation rule that uses window sequence, containment of edge fragments, and the originating analysis window, rather than treating overlap alone as evidence of duplicate detections.

### 2.5. Curated Event Records for Ecological Analysis

Existing bird-audio work provides tools for species classification and event detection, and computer-vision work provides methods for post-processing detections from overlapping windows. The measurement capability addressed here is call-type-specific event measurement in noisy field recordings, where target vocalizations can be incompletely observed, appear in multiple overlapping analysis windows, or overlap with other vocalizations and background sounds. For HLW buzz analysis, we combine event-level time-frequency localization, duplicate-window consolidation, and completeness-based curation before computing occurrence, duration, and frequency summaries. Existing public bird-audio resources and readily available pipelines do not provide this combination of annotations and outputs.

## 3. Data Collection Overview

### 3.1. Field Recording Design

AudioMoth acoustic recorders (Hill et al., 2018) were deployed near Sural village, Himachal Pradesh, India (deployment centroid: 33.1392^◦^ N, 76.4548^◦^ E; Fig. 2), within the Himalayan breeding habitat of the Hume’s Leaf Warbler. Recorders were deployed at 30 locations spanning 3,171 to 3,818 m elevation. Neighboring recorders were approximately 100 m apart and were mounted 1–3 m above ground. Recordings were collected during May–August in both 2024 and 2025, using four 15-min recording sessions every morning. The processed corpus comprised 2,573.6 hours (1,344.4 h in 2024 and 1,229.2 h in 2025). These recordings supplied the field data for model development, held-out evaluation, and ecological analysis. They contained substantial variation in background conditions, including wind, non-target bird vocalizations, and site-specific acoustic clutter. From this corpus, the finalized SpectroBird method retained 65,280 complete-buzz records for ecological analysis, including 39,105 from 2024 and 26,175 from 2025; counts across elevation bins are shown in Fig. 3.

**Figure 3:**
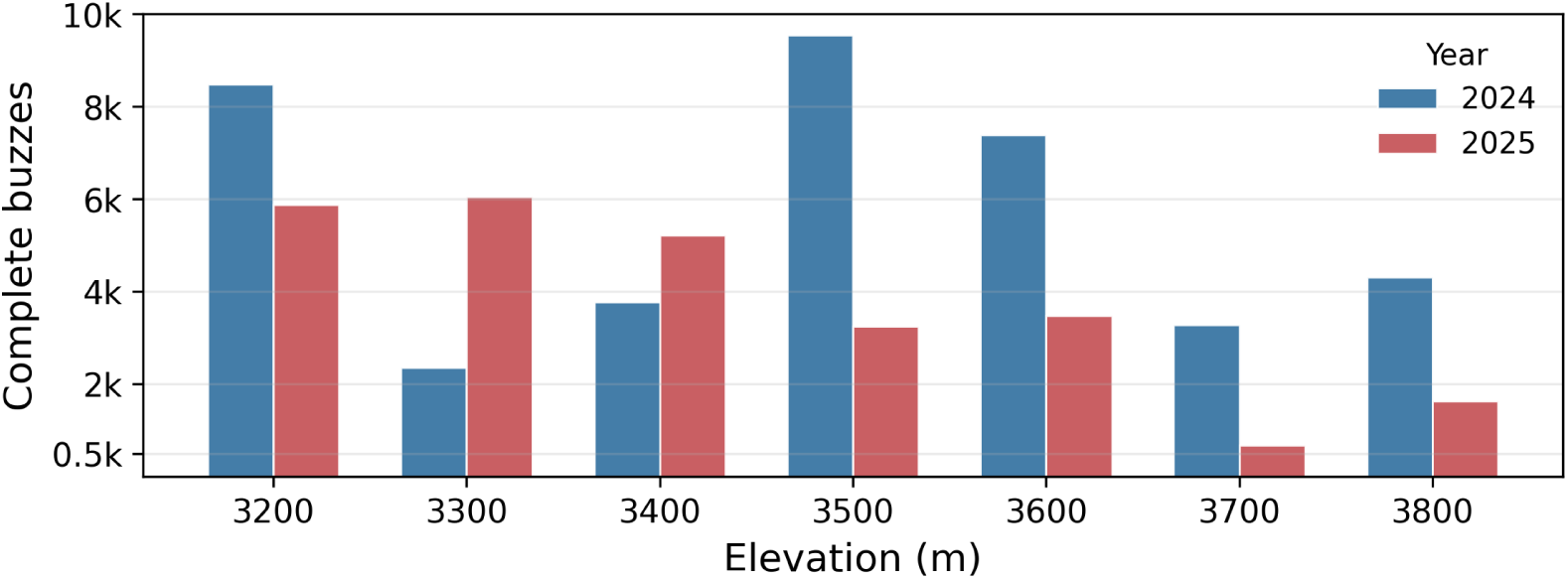
Retained complete-buzz detections in 100-m elevation bins by field season.

### 3.2. Public Audio Sources

Public recordings from Xeno-Canto and iNaturalist (Xeno-Canto Foundation, 2026; iNaturalist, 2026) are used in two roles: (i) early-stage exploration and qualitative understanding of buzz structure, and (ii) out-of-domain stress testing of generalization after model development. We treat these data separately from the held-out field test set used for primary evaluation.

Because available public resources do not provide call-type-specific time-frequency boxes or complete/incomplete event-quality labels for HLW buzzes, this study constructs both the recording corpus and annotation structure used for localization, curation, and downstream summaries.

### 3.3. Initial Data Exploration

To guide the design of the detection-localization model, an initial exploratory review was conducted using a combination of public databases and primary field recordings. All recordings were converted to spectrograms using short-time Fourier transform (STFT), and a representative subset was visually examined to assess patterns of vocal activity. This review focused on identifying features that distinguish HLW buzzes from other bird vocalizations and from background sources (e.g., low-frequency environmental sounds, wind, and other non-biological noise), in order to guide detector design. This process confirmed that HLW buzzes possess distinct, repeatable temporal and spectral features suitable for object-detection frameworks.

This analysis showed that the HLW buzz appears as a compact and visually consistent region in the spectrogram (Fig. 1b). It is typically represented by a single connected component with a smooth boundary, a shallow concavity, and an internally striped intensity pattern; this interpretation was developed with input from a coauthor with field expertise. These features make it visually distinct from the HLW call, other species’ vocalizations, and background noise. Buzzes were concentrated primarily within the 5–12 kHz frequency band, with durations typically between 1.0–1.5 s. Multiple buzzes could appear in the same frame and overlap in both time and frequency (Fig. 5).

The review also highlighted two observational regimes: fully observed buzzes, where both temporal boundaries and the internal pattern are clearly visible, and incompletely observed buzzes, where weaker portions become indistinguishable from background and only part of the event remains visible. In these incomplete cases, the apparent truncation reflects low local SNR (segments falling below the noise floor) rather than a true biological event boundary (Fig. 4).

**Figure 4:**
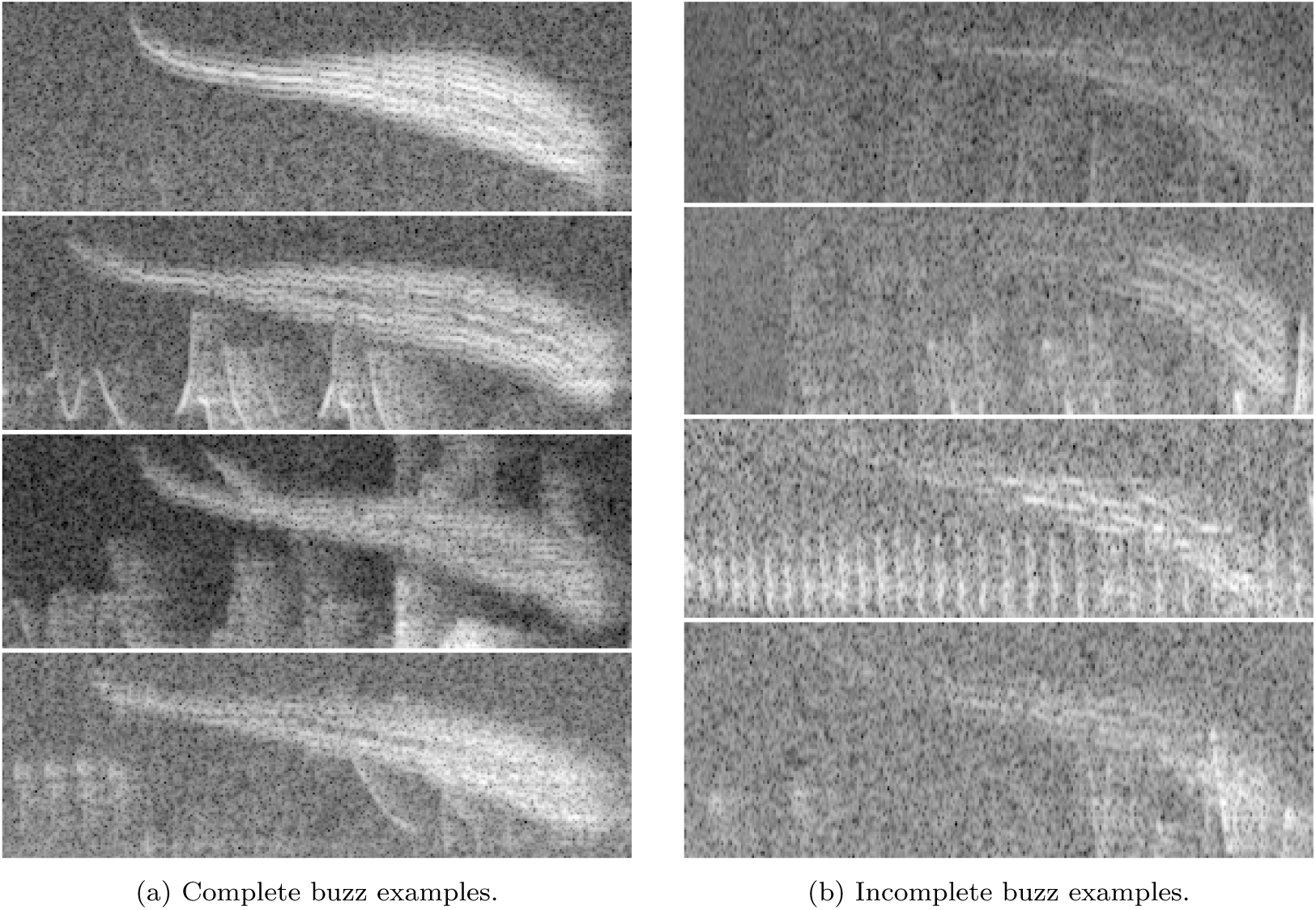
Representative complete and incomplete buzz examples. Left panel: completely observed buzzes. Right panel: incompletely observed buzzes, where portions of the signal fade into the local noise floor.

**Figure 5:**
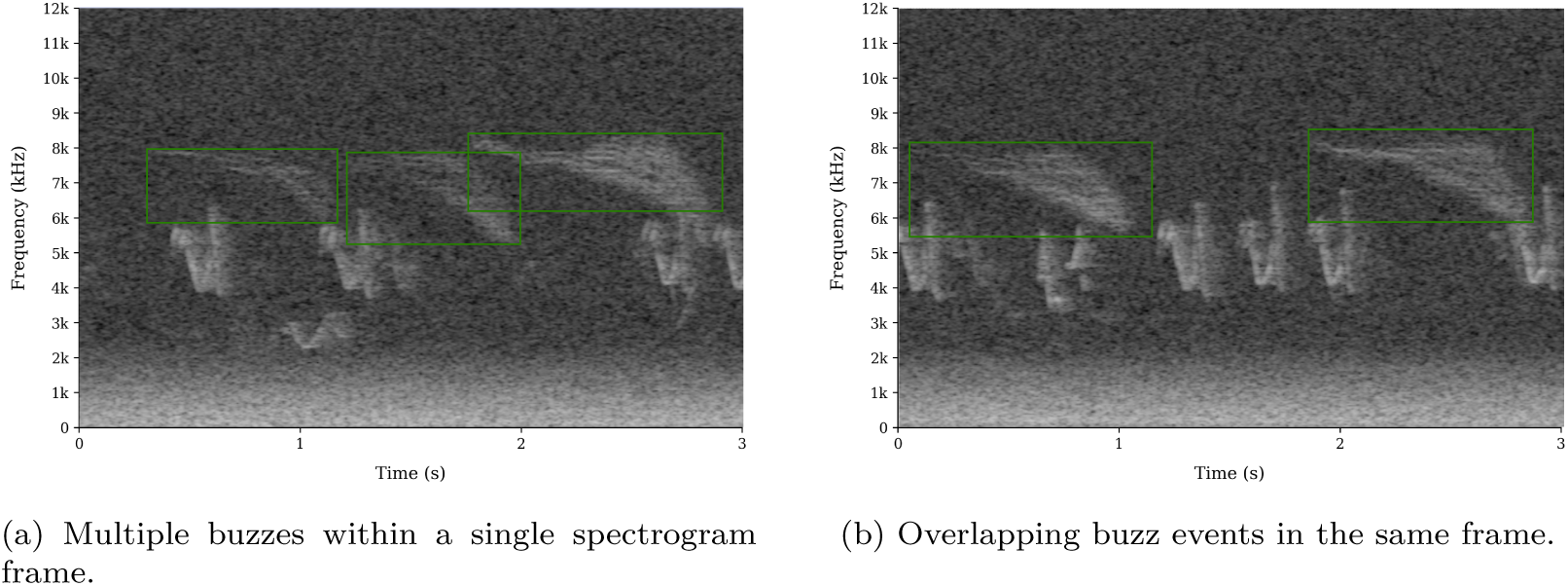
Examples of multiple and overlapping HLW buzzes.

## 4. Problem Formulation

### 4.1. Event-Level Measurement Goal

Many acoustic ecology questions require measurements tied to individual vocalizations rather than fixed recording windows. For a target call type, the relevant observation is an event: one occurrence of the vocalization that can be counted and, when complete, measured. Recovering these event records from field recordings is challenging because target events can occur under variable background noise and can overlap within the same spectrogram window (Section 3.3; Fig. 5). The computational problem is therefore to recover individual target events as the units for downstream analysis, including cases where more than one event occurs in the same analysis window.

### 4.2. Complete and Incomplete Events

Incompletely observed target events are common in field recordings: weak portions can fall below the local noise floor or be transiently masked by other sounds (Section 3.3; Fig. 4). These events may still indicate target vocal activity, but treating them as complete events can bias duration and frequency-bound summaries. The measurement task therefore requires detecting candidate target events and distinguishing events that are complete enough for structural measurement from those that are not.

### 4.3. Need for Time-Frequency Localization

Fixed-window predictions can indicate that a target vocalization is present somewhere in a clip, but they do not directly provide event identity, event boundaries, or separation of overlapping vocalizations. For analyses that depend on event counts, start/end times, duration, frequency placement, or overlap, the target must be localized in time-frequency space.

### 4.4. Need for a Task-Aligned Spectrogram Representation

Successful event-level localization also depends on how the input audio is represented. Target events may occupy restricted frequency bands or contain structure that is lost when generic representations compress, smooth, or truncate the relevant band. Log-mel representations, for example, allocate progressively fewer bins at high frequencies and can blur narrowband structure. Because HLW buzzes occupy a high-frequency band and exhibit fine horizontal striation patterns (Fig. 7), the problem requires a representation that preserves high-frequency detail in the target band.

### 4.5. Required Output Records

The required output is a set of curated event records. Each record should identify one candidate vocalization, provide start/end time and frequency bounds, carry a detection confidence, and indicate whether the event is complete enough for downstream structural analysis. For ecological applications, these event records can then be linked to recording metadata such as recorder identity, date, and site covariates.

## 5. Computational Method

### 5.1. Method Overview

The method first converts long recordings into overlapping spectrogram windows and applies a detection-localization model to identify candidate vocalizations as time-frequency boxes (Fig. 6). Because the same vocalization can appear in multiple overlapping windows, an event-consolidation step merges repeated detections while preserving genuinely overlapping vocalizations as separate events. A completeness-curation model then evaluates each consolidated event and classifies it as complete or incomplete before retained complete events are exported for downstream analysis.

**Figure 6:**
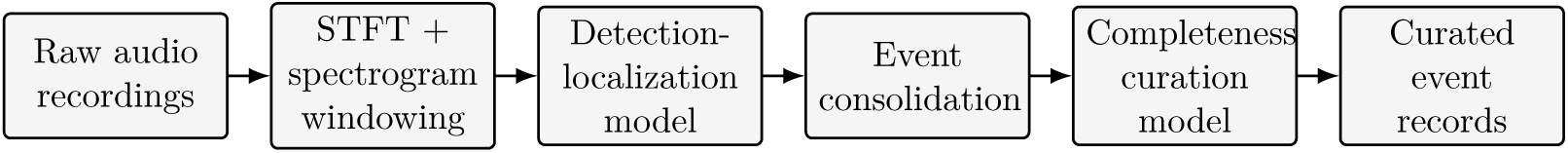
SpectroBird computational method for target-event localization, event consolidation, and completeness curation.

### 5.2. Spectrogram Representation

The representation and windowing parameters below instantiate the general method for the HLW buzz application. Audio is processed at 48 kHz using short-time Fourier transform (STFT) processing with a 1024-sample FFT, 256-sample hop length, and a linear-frequency spectrogram representation. To meet the representation requirement described in Section 4.4, detector inputs in this application use a linear frequency scale rather than a Mel scale.

BirdNET-style and standard AST spectrogram representations provide useful reference points, but they are poorly matched to the HLW buzz task. The BirdNET-style Mel representation compresses the high-frequency region where buzz structure is expressed, and the standard AST AudioSet representation assumes 16 kHz audio, placing the Nyquist frequency at 8 kHz and excluding the upper portion of many buzzes.

Figure 7 illustrates these differences on the same held-out complete buzz. The application-specific cropped linear-frequency representation retains the horizontal striation pattern visible in panel (d), whereas the comparison representations compress the high-frequency region or truncate the upper part of the buzz. The 4.125–11.625 kHz crop was chosen to comfortably bracket the observed event distribution: across consolidated field detections, the 0.001 quantile of low-frequency bounds was 4.47 kHz and the 0.999 quantile of high-frequency bounds was 10.14 kHz.

**Figure 7:**
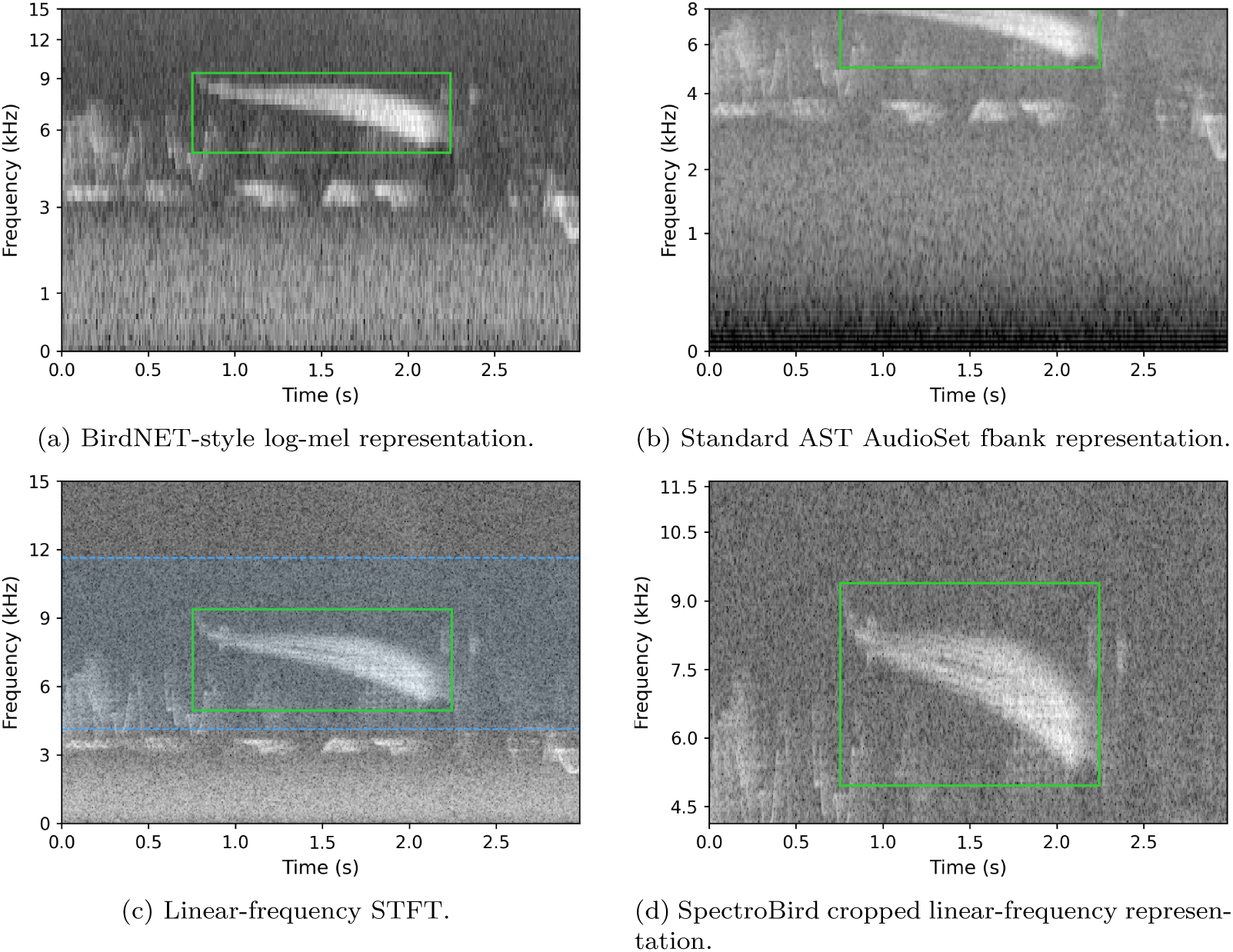
The same strong held-out HLW buzz rendered with BirdNET-style, standard AST, and linear-frequency spectrogram representations. Panel (a) shows the BirdNET-style representation, a nonlinear mel-frequency scale spanning 150 Hz–15 kHz. Panel (a) shows the standard AST AudioSet representation, a nonlinear mel-frequency scale; it assumes 16 kHz audio and therefore spans 20 Hz–8 kHz, so the upper portion of the buzz falls out of range. Panel (c) shows a linear-frequency STFT, with the SpectroBird analysis frequency band marked in blue. Panel (d) shows the final cropped 4.125–11.625 kHz linear-frequency representation used in this work. The green box marks the annotated complete buzz in all panels.

### 5.3. Analysis Windows and Detector Inputs

For this application, each analysis window spans 512 STFT frames (2.7467 s), a pragmatic choice that keeps the temporal context close to the 3 s window used in BirdNET while ensuring that detector input dimensions remain exact multiples of the YOLO stride (32). The analysis window advances by 128 frames (0.6827 s) between consecutive windows, corresponding to 75% overlap. Given this window length and stride, all observed buzzes in the 2024–2025 outputs, including the maximum-duration buzz (1.65 s), can be fully contained in at least one analysis window.

Within each window, the detector input is frequency-cropped to 4.125–11.625 kHz (spectrogram bins 88:248 in this study), flipped along the frequency axis for consistent image orientation, converted to dB relative to per-window maximum amplitude, min-max scaled to 8-bit intensity for normalization, and formatted as a 3-channel 160*×*512 image. No denoising is applied.

### 5.4. Detection-Localization Model

#### 5.4.1. Model Training

The detection-localization model is a YOLOv11 object detector (large variant, YOLOv11l) initialized from pretrained YOLOv11l detection weights and optimized on the detection-localization dataset with controlled geometric/color augmentation. The reported model was trained for 50 epochs with 160*×*512 inputs, batch size 48, initial learning rate 0.001, weight decay 0.0005, and modest brightness, translation, and scale augmentation. Additional optimizer and loss settings used the fixed YOLOv11 training configuration recorded for this run.

#### 5.4.2. Inference and Coordinate Mapping

Model predictions are converted from normalized detector-image coordinates back to absolute recording time and frequency coordinates using the analysis-window index, window stride, window duration, and fixed 4.125–11.625 kHz frequency crop. Because the input images are vertically flipped after frequency cropping, the frequency-coordinate conversion reverses the image y-axis. The explicit coordinate mapping is given in Appendix A.

### 5.5. Event Consolidation

Because analysis windows overlap by 75%, a single target vocalization can appear in several adjacent windows, including edge-truncated partial views. We therefore group repeated detections into event tracks, where a track is a group of window-level detections inferred to correspond to the same underlying event (Fig. 8a). The detections in each accepted track are consolidated into one event-level prediction (Fig. 8b).

**Figure 8:**
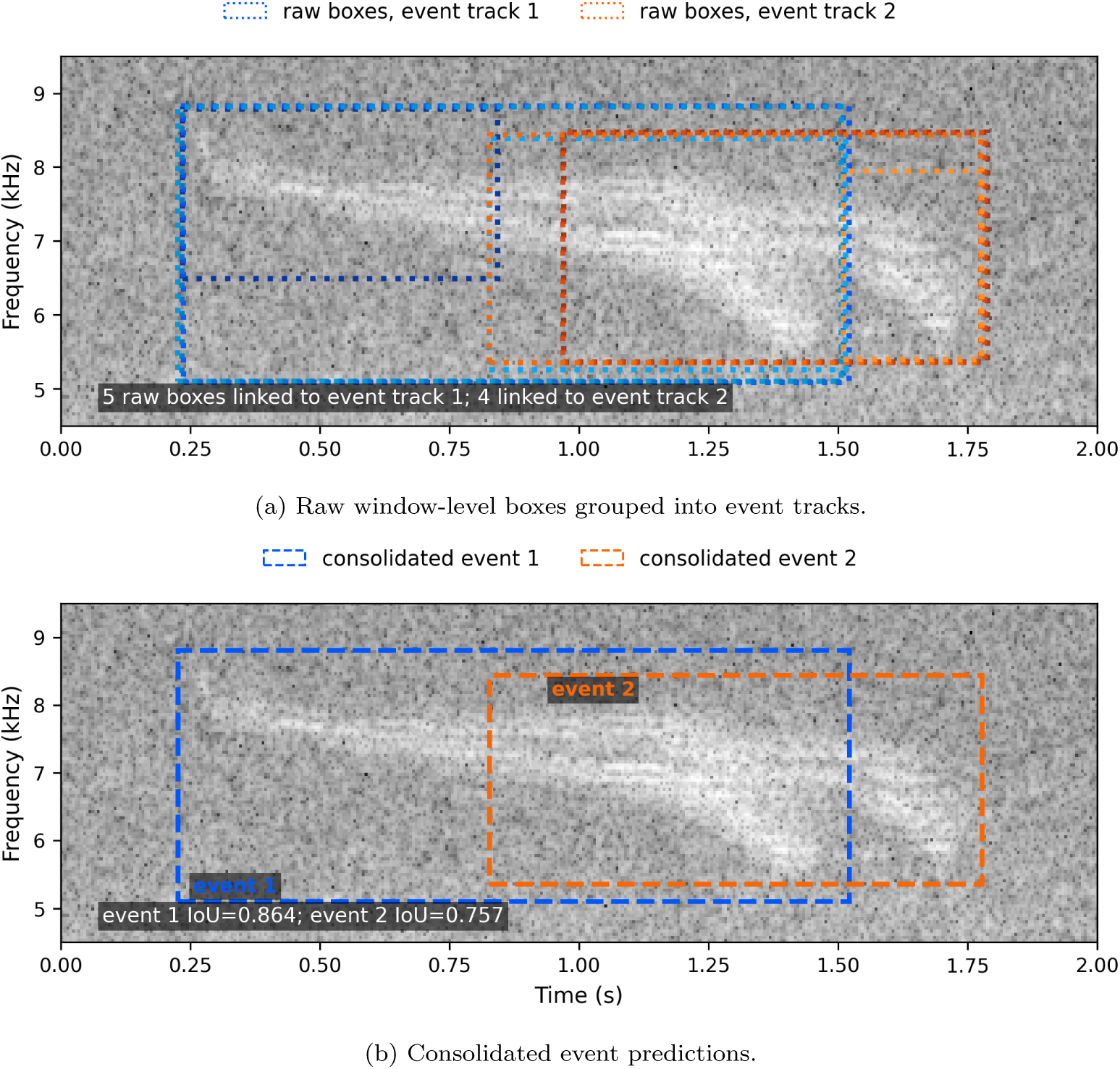
Event consolidation for overlapping buzzes. Raw window-level boxes are grouped into two event tracks, and the detections in each accepted track are consolidated into separate event-level predictions despite temporal and spectral overlap.

Consolidation is performed in absolute time-frequency coordinates. A link is a proposed association between two window-level detections that may belong to the same underlying event. Candidate same-event links are considered only between detections from nearby but distinct analysis windows within the same recording, so an event track may contain at most one detection from any given analysis window. Candidate links are scored with a fixed hand-designed association score that combines time-frequency overlap, duration and bandwidth similarity, center proximity, detector confidence, analysis-window gap, and proximity to window edges. Links that pass the fixed association thresholds are used to start new event tracks or extend existing tracks.

The procedure also handles window-edge fragments. After initial linking, a near-edge singleton detection may be absorbed into an established multi-window track if it is mostly contained by the track envelope and passes fixed score and margin gates. Detections in accepted tracks are consolidated into event-level predictions using confidence-weighted boundary fusion, with edge censoring for potentially truncated time bounds. Event confidence is the maximum member-detection confidence, and no event-confidence threshold is applied during consolidation itself; low-confidence singleton events remain available for downstream evaluation and export thresholding. Full scoring equations are provided in Appendix B, and fixed implementation parameters are summarized in Appendix Table C.1.

### 5.6. Completeness-Curation Model

The completeness-curation model performs event-quality classification on consolidated event predictions. The model estimates whether each predicted event is complete enough for structural measurement; event localization remains defined by the detection-localization model and event consolidation.

#### 5.6.1. Input Generation from Consolidated Events

At inference and export time, the completeness-curation model operates on event-level predictions after consolidation rather than raw duplicate window-level boxes. Each prediction is converted to a spectrogram crop using its time-frequency bounds and standardized to the fixed crop format used for completeness labeling (Section 6.4). The same crop-construction procedure is used for labeling and inference: crops are reconstructed from source audio using the consolidated event bounds, the predicted event is right-aligned in the standardized temporal crop, and the image is resized with preserved aspect ratio and padded to 288*×*288.

#### 5.6.2. Model Training and Validation

The completeness-curation model uses a YOLOv11l-cls image classifier initialized from ImageNet pre-trained classification weights and optimized on the complete/incomplete crop dataset. The reported model was trained for 100 epochs with 288*×*288 inputs, batch size 32, initial learning rate 0.001, weight decay 0.0005, and modest brightness, translation, and scale augmentation. Validation was run after each epoch, and the checkpoint used for inference was selected by highest validation peak F1.

#### 5.6.3. Inference and Retention Rule

At inference time, the completeness-curation model assigns a completeness confidence to each eligible event crop. Eligibility is defined by the upstream event-confidence operating threshold, so the curation model is applied only after detection-localization and event consolidation. The completeness threshold is selected on the validation split as the confidence threshold that attains peak F1, then fixed for downstream exports. Events below this threshold are labeled incomplete for export purposes and retained for diagnostic error analysis.

### 5.7. Curated Event Record Export

Curated event exports start from consolidated event predictions. For field-data summaries, retained complete-event records must pass both the validation-derived event-confidence operating threshold and the selected completeness threshold. For each retained event, the export carries event time bounds, frequency bounds, detection confidence, completeness confidence, and derived duration, frequency-center, and bandwidth fields. For field applications, these event-level records can be linked to recording metadata and study-specific ecological covariates, such as recorder identity, location, date, habitat, or elevation, to form analysis-ready tables for downstream summaries and statistics. Records with missing metadata links, non-positive duration, or invalid frequency bounds are excluded during export.

From these retained records, downstream summaries can be computed at both the event level and at user-defined sampling units such as recorders, recorder-years, sites, or days. For sampling-unit-level summaries, retained events are first averaged within the chosen unit, and covariate-level summaries are then computed across unit-level means rather than individual events. This prevents high-activity sampling units from dominating summaries of duration or frequency structure.

## 6. Dataset Development

### 6.1. Dataset Splits and Evaluation Subsets

Dataset construction uses two data granularities: analysis windows for the detection-localization model and event crops for the completeness-curation model. We define five primary train/validation/test splits and one additional disjoint public-audio evaluation subset used for generalization analysis:

- *D*_det-train_, *D*_det-val_: detection-localization training and validation sets.
- *D*_cur-train_, *D*_cur-val_: completeness-curation training and validation sets.
- *D*_test_: held-out, manually annotated field recordings for model comparison.
- *D*_public_: a disjoint public-audio holdout used for additional generalization analysis, consisting of public-recording submissions unrelated to those used in early training-data development.

The relationships among these partitions and their construction paths, along with the associated dataset counts, are summarized in Fig. 9 and Table 1.

**Figure 9:**
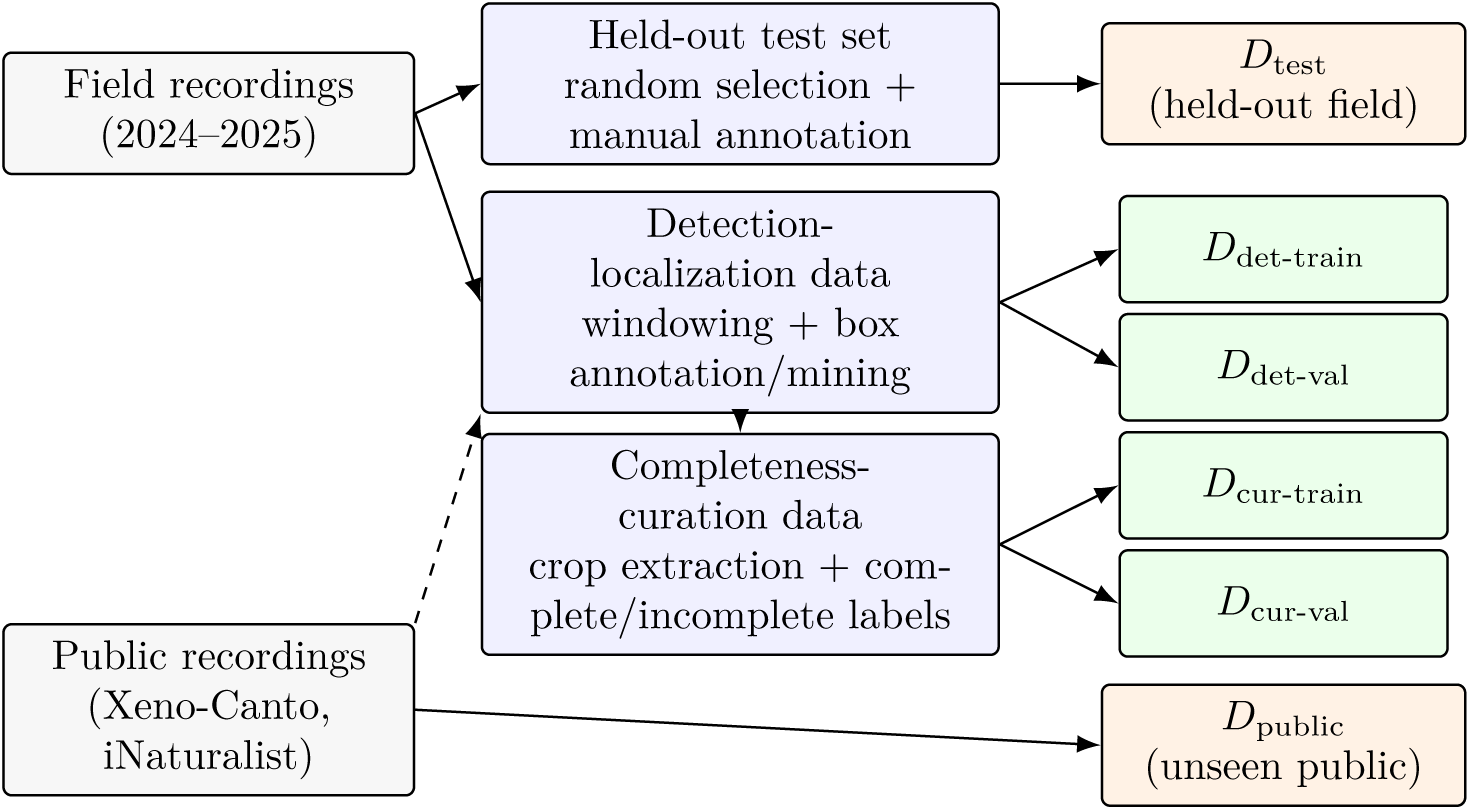
Data flow and partitioning used in this study. Solid arrows indicate primary construction paths. Dashed arrow indicates limited use of other public submissions in early seed curation. *D*_public_ denotes a disjoint public-audio holdout and was not itself used for training, model selection, or seed curation.

**Table 1:** Dataset counts for major training and evaluation sets used by the SpectroBird method.

| Component | Split/Class | Count |
| --- | --- | --- |
| Detection-localization | training windows | 34,385 |
| Detection-localization | validation windows | 1,799 |
| Detection-localization | training buzz boxes | 13,718 |
| Detection-localization | validation buzz boxes | 716 |
| Completeness-curation | train complete | 17,283 |
| Completeness-curation | train incomplete | 12,234 |
| Completeness-curation | val complete | 506 |
| Completeness-curation | val incomplete | 336 |
| Held-out field test | recordings | 54 |
| Held-out field test | final annotated buzz events | 1,081 |
| Held-out field test | final complete buzz events | 391 |
| Held-out field test | final incomplete buzz events | 690 |
| Unseen public audio | files analyzed | 38 |
| Unseen public audio | buzz vocalizations | 125 |

### 6.2. Test-Set Annotation Summary

The primary held-out test set contains 54 field recordings (15 minutes each; 13.5 h total), selected at random from the 2024 field recordings, with 1,081 final HLW buzz-event boxes. Events were labeled in Audacity (Audacity Team, 2026) as time-frequency boxes (start/end time and lower/upper frequency), with non-buzz audio left unlabeled. Each file received two independent researcher annotation passes, followed by an expert adjudication pass to finalize ground-truth labels used only for held-out evaluation. This expert-adjudicated held-out set supports detection-localization evaluation.

Complete/incomplete labels used for detection-localization performance evaluation follow the same expert-guided visibility criteria used for completeness-curation crop labeling, defined below. However, completeness-curation performance is evaluated only on its internal validation split rather than on an independently expert-adjudicated test set. Timeline-level inter-annotator agreement, quantified with Krippendorff’s *α* over time bins including silence intervals, was 0.92. Detailed adjudication mechanics, including weighted-vote logic and agreement implementation details, are documented in Appendix D. Review of preliminary detector outputs prompted a final annotation-QC pass focused on duplicate or misplaced boxes, overlapping-event separation, and boundary placement. This step was important for fair comparison across all detection models because the same reference events determine window-level labels, stratum membership, precision-recall accounting, and direct event-target matching. The final held-out event labels contain 391 complete buzzes and 690 incomplete buzzes; analyses run before this pass were used only for error analysis.

### 6.3. Training Data Curation: Buzz Event Localization

Training data were built iteratively. A small initial dataset combining examples from Xeno-Canto, iNaturalist, and field recordings was manually annotated in CVAT.ai (CVAT.ai, 2026) using time-frequency bounding boxes around buzz events, and used to train an initial detector. That model then mined candidate positives/negatives from larger unlabeled field audio. Candidate windows were sampled across detector-confidence strata and elevation bands from non-test recordings, then manually verified and box-corrected in CVAT. Recurrent false-positive patterns were explicitly retained as hard negatives. Repeating this loop expanded representation of background conditions, overlapping events, non-stationary noise, and acoustically similar hard negatives.

During manual box annotation and correction, window-edge buzzes were labeled only when the visible fragment remained recognizable as part of a buzz. In those cases, the box marked the visible fragment within the analysis window; severely truncated or ambiguous fragments were not annotated as target boxes.

For classifier baseline comparisons, a clip-level binary dataset was also derived from detection-localization box annotations using the window-label construction protocol defined in Section 7.3; the resulting benchmark comparison is described in Section 7.6.2.

### 6.4. Training Data Curation: Buzz Crop Completeness Labeling

Completeness curation uses a binary crop-level label scheme: *complete* versus *incomplete*. To build this corpus, candidate buzz crops are generated from non-test field detections. Here, each crop is the time-frequency rectangular region of the spectrogram corresponding to a predicted buzz box. Each detection is converted to a standardized crop with a fixed temporal width of 288 frames, with the predicted event right-aligned in the crop and left-side context or padding added as needed. Temporal padding is sampled from local non-buzz regions to preserve realistic background texture. The time dimension is preserved at 288 frames, and the frequency dimension is padded with gray background values to form 288*×*288 classifier inputs.

Completeness-curation crops were uploaded to CVAT.ai for manual binary annotation. The annotation rules were developed with guidance from a coauthor with field expertise, and annotated crops were then organized into train/validation splits (counts in Table 1). A crop is labeled *complete* only when both conditions hold:

- temporal containment: both start and end are visible within the crop;
- continuous structure: the buzz appears as one continuous structure rather than an edge-truncated fragment.

Crops that fail either condition are labeled *incomplete*. Representative labeled crop examples are shown in Fig. 4.

## 7. Model Evaluation

### 7.1. Evaluation Claims

The evaluation tests four claims about the SpectroBird method. First, consolidated detector outputs recover annotated buzz events and their time-frequency boundaries on held-out field recordings. Second, under a shared fixed-window benchmark, the detector remains competitive with strong window classifiers while retaining event-level outputs. Third, the curation model distinguishes complete from incomplete event crops well enough to support complete-event export. Fourth, the finalized method transfers to unseen public recordings from outside the field corpus.

### 7.2. Evaluation Sets

We evaluate models on a held-out, manually annotated field test set and a separate unseen public-audio set. Training and mining are performed on non-test recordings. Test recordings are excluded by filename during mining and inference to prevent data leakage. Thresholds used for downstream exports and retained-event decisions are selected on development/validation data. Cross-model benchmark comparisons use average precision (AP) as the primary summary metric; peak F1 from test-set threshold sweeps is retained only as a secondary diagnostic and is not interpreted as a deployment operating threshold.

From the held-out field test set, we construct three reporting strata for the detection-localization evaluation: (i) all buzzes + background, (ii) complete buzzes + background, and (iii) incomplete buzzes + background. Primary cross-model benchmark reporting focuses on the all-buzz and complete-buzz strata, because structural measurements such as duration and frequency extent require completely observed events. In stratum-restricted analyses, the opposite buzz stratum is treated as ignore rather than background, while true background negatives are retained. This avoids penalizing a model for detecting a real buzz that falls outside the reporting stratum, and lets the complete-buzz and incomplete-buzz rows isolate retrieval within each event-quality class. All-buzz performance quantifies general buzz retrieval under field conditions, while complete-buzz performance isolates retrieval of events eligible for structural and ecological summaries. The stratum of incomplete buzzes remains useful as a secondary diagnostic for distant, occluded, or otherwise low-SNR events, but is not emphasized in the primary benchmark table.

### 7.3. Shared Window-Level Benchmark Protocol

For the shared classifier benchmark, each test recording is divided into the same 2.7467 s analysis windows with a 0.6827 s stride. Each model contributes one confidence score per window. The classifier baselines already produce one window score directly; for the SpectroBird detector, we use the highest detection-box confidence within the window for this benchmark only. This conversion discards localization information, which is evaluated separately in the direct event-level analyses.

Window labels are assigned separately for each reporting stratum: all buzzes, complete buzzes, or incomplete buzzes. We report two complementary window-labeling criteria. Under the *complete-within-window* criterion, a window is positive if it fully contains at least one reference event from the reporting stratum. Under the *minimum-overlap* criterion, a window is positive if it overlaps a reference event from the reporting stratum by at least 0.2 s. This threshold was selected during analysis review because buzz structure begins to become visually identifiable at approximately this amount of temporal overlap. Windows with no overlap with any reference event in the reporting universe are negatives. Windows that overlap a reference event but do not satisfy the selected positive criterion are treated as *ignore* and excluded from TP/FP/FN accounting. The complete-within-window criterion emphasizes complete-event availability within a frame, whereas the minimum-overlap criterion tests model behavior on boundary windows where only part of the event is visible.

We implement scoring with the following steps:

1. **Construct common evaluation windows.** Enumerate all quarter-step windows across test recordings and assign each model a score per window (missing model outputs are set to 0).
2. **Assign positives, negatives, and ignore windows.** Apply the selected window-labeling criterion for positives. Mark non-qualifying overlap windows as ignore; keep non-overlap windows as negatives.
3. **Apply stratum masks.** For complete-only and incomplete-only strata, treat the opposite buzz stratum as ignore and exclude from scoring.
4. **Threshold sweep and confusion accounting.** For each confidence threshold, classify non-ignored windows by score, compute TP/FP/FN, and record precision, recall, and F1.

Precision, recall, and F1 are computed at each threshold from the resulting TP/FP/FN counts.

### 7.4. Event Matching and Confidence Thresholding

Direct event-level evaluation compares consolidated SpectroBird predictions with the final held-out time-frequency boxes. Matching is performed independently within each recording using one-to-one assignment between eligible predictions and reference events. The assignment score combines time-frequency overlap, time overlap, frequency overlap, smaller-box coverage, center proximity, and prediction confidence. IoU is reported after matching rather than used as a hard event-identity gate, so plausible same-event matches with imperfect boundaries remain available for localization-error summaries.

For event-level precision-recall sweeps, predictions below the current event-confidence threshold are removed and one-to-one assignment is rerun. Matched predictions are true positives, unmatched eligible predictions are false positives, and unmatched reference events are false negatives. For complete-only or incomplete-only reporting, the opposite buzz stratum is treated as ignore. Full event-matching parameters are provided in Appendix Table C.2.

### 7.5. Metrics

Cross-model comparison uses the shared window-level protocol in Section 7.3. We sweep confidence to generate precision-recall curves and use AP as the primary model-comparison metric. For the main benchmark table, uncertainty is summarized with paired 95% recording-level bootstrap confidence intervals for the AP difference between the SpectroBird detector and each classifier baseline, preserving dependence among overlapping windows from the same recording. Peak F1 from the same threshold sweeps is treated only as a secondary descriptive summary.

For direct event-level evaluation, we report event-retrieval AP, peak F1, precision, recall, TP/FP/FN counts, median two-dimensional IoU, and boundary-error summaries for matched true-positive events. These diagnostics evaluate the boundary geometry needed for downstream duration and frequency summaries rather than window-level buzz presence alone.

The completeness-curation model is evaluated as a binary classifier on complete versus incomplete labels using its validation split. This split is used for internal model selection and threshold calibration rather than as a separate expert-adjudicated test set. We report validation precision, recall, F1, and area under the PR curve (AUCPR). The completeness-curation check-point is selected by highest validation peak F1. The completeness threshold is then set to the confidence threshold that attains peak F1 for that selected checkpoint and fixed for downstream duration analysis.

### 7.6. Results

#### 7.6.1. Detection-Localization Event Retrieval and Boundary Accuracy

Direct event-level evaluation measures how well consolidated predictions recover individual annotated buzz events and their time-frequency boundaries on the final held-out event annotations.

The detection-localization model performed strongest on complete buzzes, the stratum most directly relevant to downstream duration exports (Table 2). For complete buzzes, AP was 0.9995 and peak F1 was 0.9911; at the peak-F1 threshold, the model recovered 388 of 391 complete buzzes with four false positives. Performance remained strong for all buzzes combined (AP = 0.9600, peak F1 = 0.9299), while incomplete-buzz performance was lower, consistent with the greater ambiguity of distant, truncated, masked, or otherwise low-SNR events.

**Table 2:**
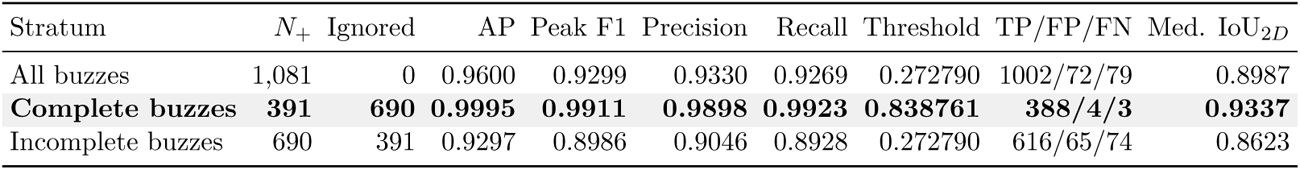
Direct event-level performance on the final held-out test set after sequence-aware event consolidation and one-to-one event-identity matching. Peak F1 and the threshold column are reported from a full event-confidence threshold sweep as post hoc diagnostic summaries, not as deployment-threshold estimates. For stratum-restricted rows, the opposite buzz stratum is treated as an ignore set. The shaded row indicates the evaluation stratum most directly relevant to downstream duration exports; the field-export confidence gate is selected separately from validation data.

| Stratum | $N_+$ | Ignored | AP | Peak F1 | Precision | Recall | Threshold | TP/FP/FN | Med. IoU<sub>2D</sub> |
| --- | --- | --- | --- | --- | --- | --- | --- | --- | --- |
| All buzzes | 1,081 | 0 | 0.9600 | 0.9299 | 0.9330 | 0.9269 | 0.272790 | 1002/72/79 | 0.8987 |
| **Complete buzzes** | **391** | **690** | **0.9995** | **0.9911** | **0.9898** | **0.9923** | **0.838761** | **388/4/3** | **0.9337** |
| Incomplete buzzes | 690 | 391 | 0.9297 | 0.8986 | 0.9046 | 0.8928 | 0.272790 | 616/65/74 | 0.8623 |

Matched true-positive events also showed high boundary accuracy (Table 3). For complete buzzes, the median IoU_2*D*_ was 0.934 and the median absolute duration error was 0.0202 s. Localization was stronger for complete buzzes than incomplete buzzes, and full localization-error distributions are provided in Appendix Fig. F.1. Representative held-out predictions are shown in Fig. 10, including a long complete buzz, overlapping buzzes, background bird vocalizations, and crowded sequences. Representative false-positive and false-negative cases are provided in Appendix Fig. E.1.

**Figure 10:**
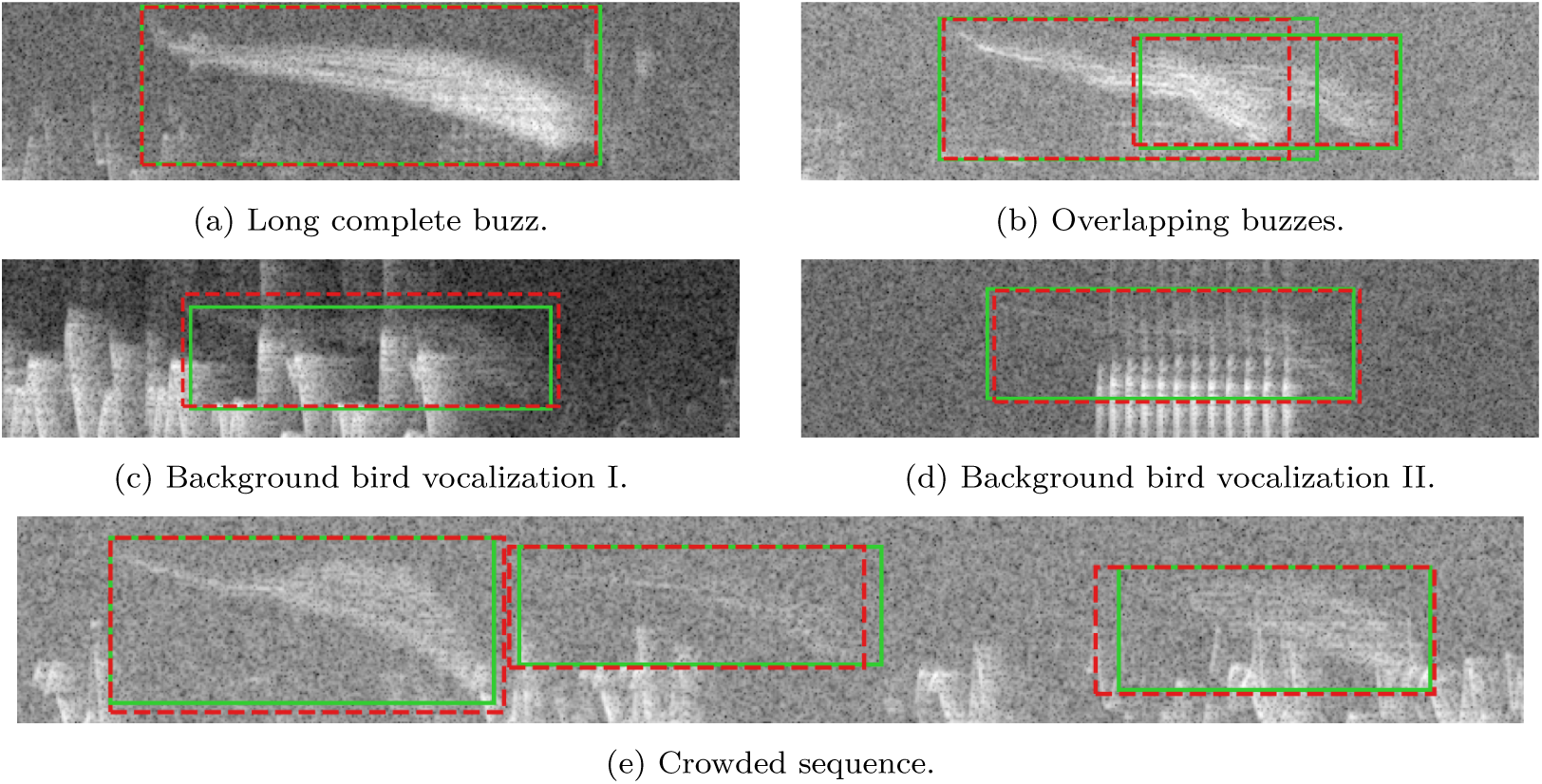
Held-out detection-localization examples after overlapping-window consolidation and one-to-one event matching. Solid green boxes mark reference events and dashed red boxes mark matched consolidated predictions. In all panels, time increases from left to right and frequency from bottom to top. Panels illustrate a long complete buzz, overlapping buzzes resolved as separate events, detections amid background bird vocalizations, and a crowded sequence of nearby buzzes.

**Table 3:** Detection-localization error for matched true-positive events at the all-buzz peak-F1 event-confidence threshold. Temporal errors are median absolute errors in seconds; frequency errors are median absolute errors in Hz; IoU median and IQR are reported separately.

| Metric | All buzzes | Complete buzzes | Incomplete buzzes |
| --- | --- | --- | --- |
| Matched TP | 1002 | 391 | 611 |
| IoU<sub>2D</sub> median | 0.899 | 0.934 | 0.863 |
| IoU<sub>2D</sub> IQR | 0.824–0.941 | 0.900–0.958 | 0.776–0.918 |
| Start time (s) | 0.0178 | 0.0091 | 0.0334 |
| End time (s) | 0.0181 | 0.0139 | 0.0209 |
| Duration (s) | 0.0328 | 0.0202 | 0.0513 |
| Low freq. (Hz) | 80.6 | 62.8 | 93.8 |
| High freq. (Hz) | 67.5 | 59.6 | 79.7 |
| Center freq. (Hz) | 57.9 | 44.5 | 66.0 |

#### 7.6.2. Comparison with Window-Level Classifiers

A central evaluation question is whether the detector can match strong window-level classifiers on detection quality while also providing event-localization outputs. We therefore compare the SpectroBird detector directly against two baselines trained on the same analysis windows:

- **AST benchmark**: the Audio Spectrogram Transformer, used here as a benchmark audio detection model on full analysis windows.
- **YOLOv11-cls baseline**: a YOLOv11 image-classification variant adapted to full-window buzz-presence detection.

For comparability with the detector inputs, AST was trained on the same 2.7467 s analysis windows and the same 4.125–11.625 kHz linear-frequency spectrogram representation used by SpectroBird, rather than the standard AudioSet AST spectrogram representation. This departs from standard AST preprocessing, which assumes 16 kHz audio and a log-mel representation; in our setting, that representation truncates most buzzes above 8 kHz and compresses the remaining high-frequency structure. Preliminary experiments with the standard AST representation performed poorly, so we used the task-aligned representation for the benchmark comparison. Accordingly, the AST benchmark should be interpreted as an AST-family classifier trained on our high-band linear-frequency representation, not as the standard AudioSet AST representation. Both classifiers output a single confidence score *p*(buzz present | window) per window. All three models are trained and evaluated under the same split constraints and shared window-level protocol in Section 7.3, ensuring a fair formulation-level comparison.

Table 4 provides a controlled comparison asking whether the richer detection-localization formulation sacrifices basic buzz-retrieval performance when all three models are evaluated under the same coarse window-level protocol in Section 7.3. AST and the YOLOv11-cls baseline use their direct window confidence scores, whereas the SpectroBird detector uses the maximum detection-box confidence in each window. We report AP under the two window-labeling criteria defined there: fully within window containment and partial within-window overlap (*d*_match_ = 0.2 s).

**Table 4:** Primary model comparison on the manually annotated test set using average precision (AP). Background negatives are retained in all reported strata. Bracketed intervals denote paired 95% recording-level bootstrap confidence intervals for the SpectroBird detector relative to each classifier baseline, using resampling over the 54 held-out recordings. FW = fully within window; PW = partially within window (minimum-overlap criterion with *d*_match_ = 0.2 s).

| Context | Detector AP | $\Delta$ vs AST | $\Delta$ vs YOLOv11-cls |
| --- | --- | --- | --- |
| PW \| All | 0.9234 | 0.0049 [−0.012, 0.016] | −0.0227 [−0.049, −0.011] |
| PW \| Complete | 0.9874 | 0.0178 [0.008, 0.031] | 0.0001 [−0.006, 0.003] |
| FW \| All | 0.9594 | −0.0219 [−0.046, −0.010] | −0.0262 [−0.055, −0.011] |
| FW \| Complete | 0.9984 | −0.0015 [−0.009, 0.000] | −0.0015 [−0.009, 0.000] |

Under the fully within-window criterion, the SpectroBird detector is modestly lower than both classifier baselines for all buzzes, while complete-buzz AP is near ceiling for all three models. Under the partial within-window criterion, the detector and AST are similar for all buzzes, whereas the YOLOv11-cls baseline is somewhat higher overall. For complete buzzes, the setting most relevant to partially visible but fully observed events, the detector exceeds AST and is effectively tied with the YOLOv11-cls baseline. Taken together, these comparisons show that the detector remains competitive under a shared window-level benchmark while retaining event timing and frequency outputs.

#### 7.6.3. Completeness-Curation Results

This subsection reports complete/incomplete classification quality for the completeness-curation model on its held-out validation split, separate from detection performance. This validation set was developed with expert guidance, and the metrics below evaluate the selected classifier within that validation framework rather than on a separate external adjudication set. Metrics are reported for the selected checkpoint at the validation peak-F1 threshold.

Representative labeled crops are shown in Fig. 4, illustrating the visual structure of both “complete” and “incomplete” examples used in completeness-curation evaluation.

Using the selected checkpoint and the peak-F1 validation threshold *θ*_cur_ = 0.530 on *p*(complete), the completeness-curation model achieved precision 0.939, recall 0.941, F1 0.940, and AUCPR 0.984 on *D*_cur-val_ (842 validation samples: 506 complete, 336 incomplete). At *θ*_cur_, the confusion counts were 476 / 31 / 30 / 305 (TP / FP / FN / TN), corresponding to accuracy 0.928.

#### 7.6.4. Generalization on Unseen Public Audio

An important question is whether the finalized method transfers beyond the field-recording conditions and curated development data used in the main study. We therefore evaluate it on unseen public recordings as an out-of-domain generalization check, using different Xeno-Canto submissions from those involved in early training-data development and spanning different recordists, devices, and locations. Figure 11 provides curated qualitative examples, and the quantitative summary below focuses on whether the method can recover completely observed buzzes under these heterogeneous public-audio conditions. We evaluated the detection-localization model on 38 Xeno-Canto clips containing 125 annotated buzz targets.

**Figure 11:**
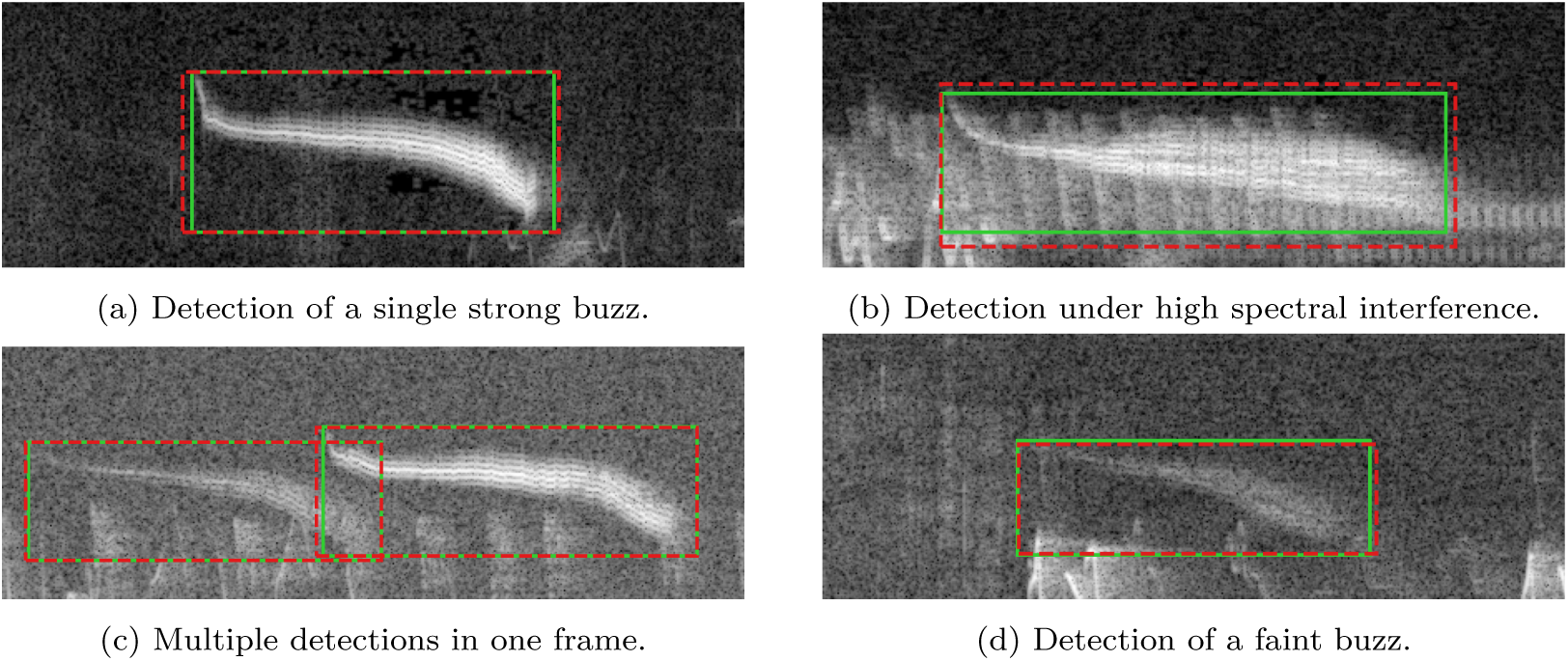
Representative SpectroBird detections on unseen Xeno-Canto recordings. Solid green boxes show annotated references and dashed red boxes show consolidated event predictions; horizontal axis is time (s).

We focus here on the completely observed subset defined on the public targets. In these unseen public recordings, the method recovered all 69 completely observed buzzes (69/69; recall = 1.000). Review at the summary threshold *θ* = 0.805 found no spurious off-target buzz localizations; the remaining four excess event counts reflected duplicate representations of already-detected buzzes.

## 8. Ecological Results

Applying SpectroBird to the HLW field recordings produced recorder-linked complete-buzz records for ecological analysis. We first used these records to test our two elevation predictions. The duration prediction was supported: complete buzzes were longer at lower elevations. The occurrence prediction was not supported: complete-buzz occurrence showed no corresponding elevational decline. We additionally examined differences in complete-buzz occurrence between years and the similarity of day-of-year occurrence patterns. Unless otherwise noted, the simple ecological models below use recorder-year summaries as observations rather than individual buzz events.

### 8.1. Exported Complete-Buzz Records

Across 2024 and 2025, SpectroBird produced 200,737 event-level predictions after consolidation (134,701 in 2024; 66,036 in 2025). Applying the validation-derived detection operating threshold (*θ*_det_ = 0.539), completeness curation, and export data-cleanliness checks for metadata linkage and valid bounds retained 65,280 complete-buzz records for ecological analysis (39,105 in 2024; 26,175 in 2025). This validation-derived export gate is separate from the post hoc test-set threshold sweeps reported in Table 2. All occurrence and duration summaries below use this retained complete-buzz set.

### 8.2. Buzz Occurrence Across Elevation

For occurrence, retained complete-buzz records were summarized at the recorder-year level as complete buzzes per recorded hour, including eligible recordings with zero retained events. Because occurrence was calculated from consolidated event records rather than positive windows, nearby or overlapping buzzes contributed as separate event counts (Fig. 12). Contrary to the prediction that buzzes would be more frequent at lower elevations, recorder-level occurrence did not show a clear monotonic elevational trend. In simple linear models of complete buzzes per hour versus elevation, slopes per 100 m were weak in 2024 (2.3 *±* 2.8, *t*-statistic = 0.82, *p*-value = 0.419), 2025 (−2.5 *±* 2.6, *t*-statistic = −0.95, *p*-value = 0.349), and pooled recorder-years (−0.2 *±* 1.9, *t*-statistic = −0.12, *p*-value = 0.902). Thus, the predicted decrease in occurrence with elevation was not supported.

**Figure 12:**
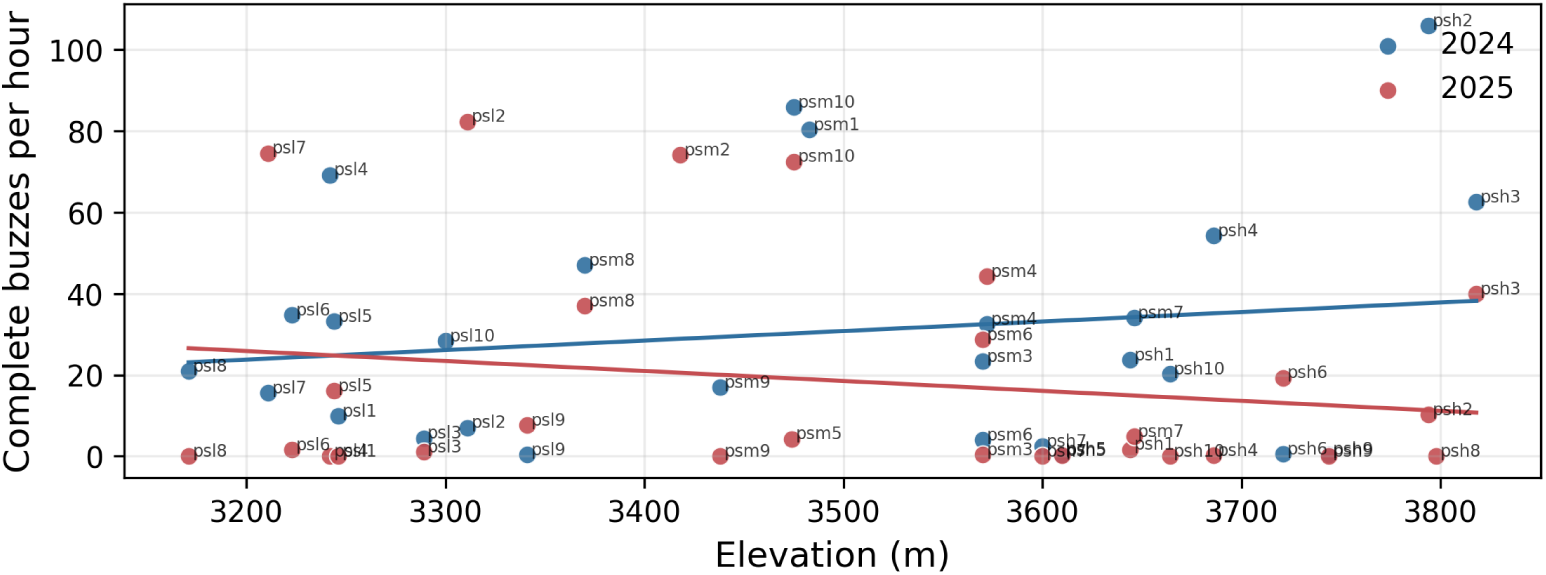
Complete-buzz occurrence versus elevation. Occurrence is summarized as retained complete buzzes per recorded hour for each recorder-year.

### 8.3. Buzz Duration Across Elevation

Duration analyses used the retained complete-buzz event set, because incomplete detections can underestimate event duration. Retained complete events were linked to recorder metadata and summarized at the recorder-year level before elevation analysis (Fig. 13). In contrast to occurrence, mean complete-buzz duration declined with elevation. In simple linear models of recorder-level mean duration versus elevation, the estimated slope was weakly negative in 2024 (−10.5 *±* 6.1 ms per 100 m, *t*-statistic = −1.72, *p*-value = 0.097), stronger in 2025 (−27.1 *±* 9.9 ms per 100 m, *t*-statistic = −2.73, *p*-value = 0.013), and negative in pooled recorder-years (−17.9 *±* 5.6 ms per 100 m, *t*-statistic = −3.20, *p*-value = 0.0024). Thus, the duration prediction was supported: complete buzzes were longer at lower elevations.

**Figure 13:**
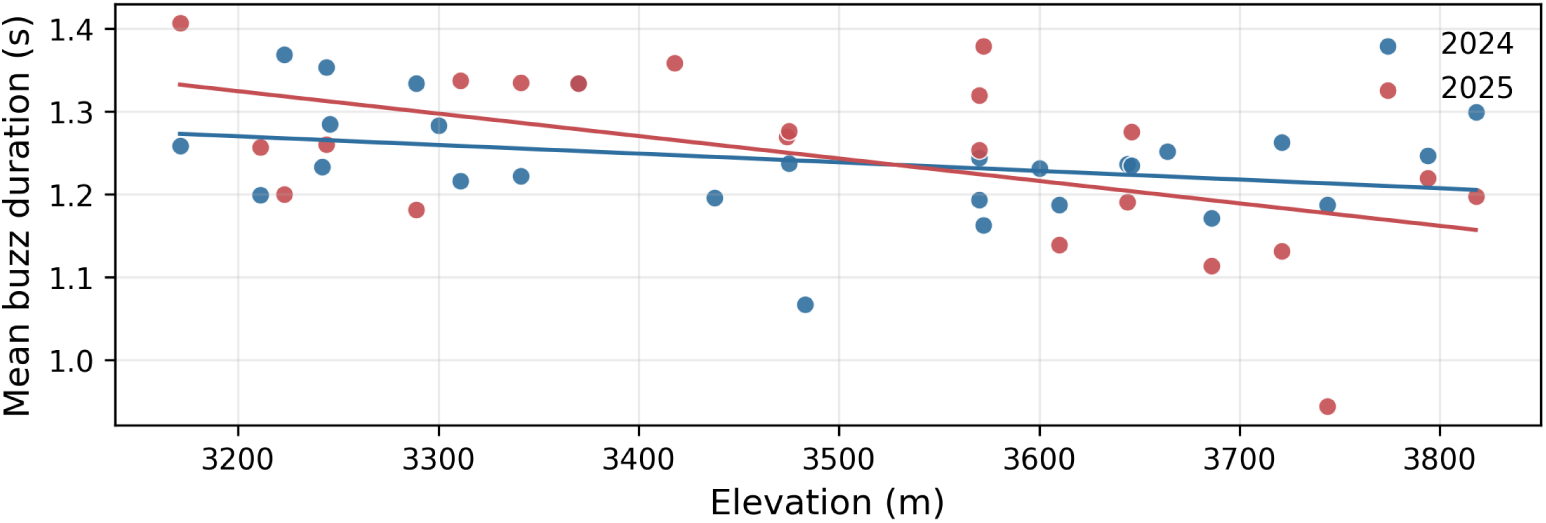
Complete-buzz duration versus elevation from recorder-linked event records. Duration is summarized as recorder-year mean duration of retained complete buzzes.

As a sanity check on the duration–elevation result, we examined detection-localization duration error on the held-out complete-buzz test set by elevation band. Complete-buzz recovery was near ceiling across bands (low: 49/49; mid: 185/187; high: 154/155), and median absolute duration error was small relative to the ecological contrast: approximately 15–27 ms across elevation bands, compared with an observed low-to-high recorder-level mean duration difference of about 80–90 ms. Signed error also did not indicate a detector bias that could explain longer buzzes at lower elevations; low- and mid-elevation durations were slightly underestimated, while high-elevation durations were slightly overestimated. Thus, held-out localization errors were smaller than the observed duration contrast and would tend to attenuate, rather than produce, the observed decline in complete-buzz duration with elevation.

For the elevation summaries, recorder is the primary descriptive unit: retained events are averaged within recorder, and recorder-level means are then summarized by elevation band. This reflects how buzz duration versus elevation would be analyzed downstream, because events from the same recorder are not independent observations of elevation. Event-level summaries still describe the exported event population, but recorder-level summaries reduce pseudo-replication and prevent high-activity recorders from dominating apparent elevation patterns.

### 8.4. Between-Year and Seasonal Occurrence Patterns

Overall retained complete-buzz occurrence was descriptively lower in 2025 (26,175 complete buzzes; 21.3 h^−1^ over 1,229.2 recorded hours) than in 2024 (39,105 complete buzzes; 29.1 h^−1^ over 1,344.4 recorded hours; Fig. 14a). In a simple model including year and elevation, however, the 2025 term was not statistically significant (−11.7 *±* 7.7 complete buzzes h^−1^, *t*-statistic = −1.53, *p*-value = 0.133), and the elevation term was also not statistically significant (*p*-value = 0.947). Thus, the aggregate summaries show lower complete-buzz occurrence in 2025, but the adjusted analysis does not establish a year effect. The relative day-of-year occurrence patterns were strongly correlated between years (Fig. 14b). After each year’s smoothed occurrence curve was scaled to its own maximum, the curves were strongly aligned across the full 71-day shared portion (Pearson *r* = 0.843; Spearman *ρ* = 0.856). Because the curves are built from retained consolidated events, close or overlapping buzzes contribute as separate vocalizations rather than being collapsed into window-level presence.

**Figure 14:**
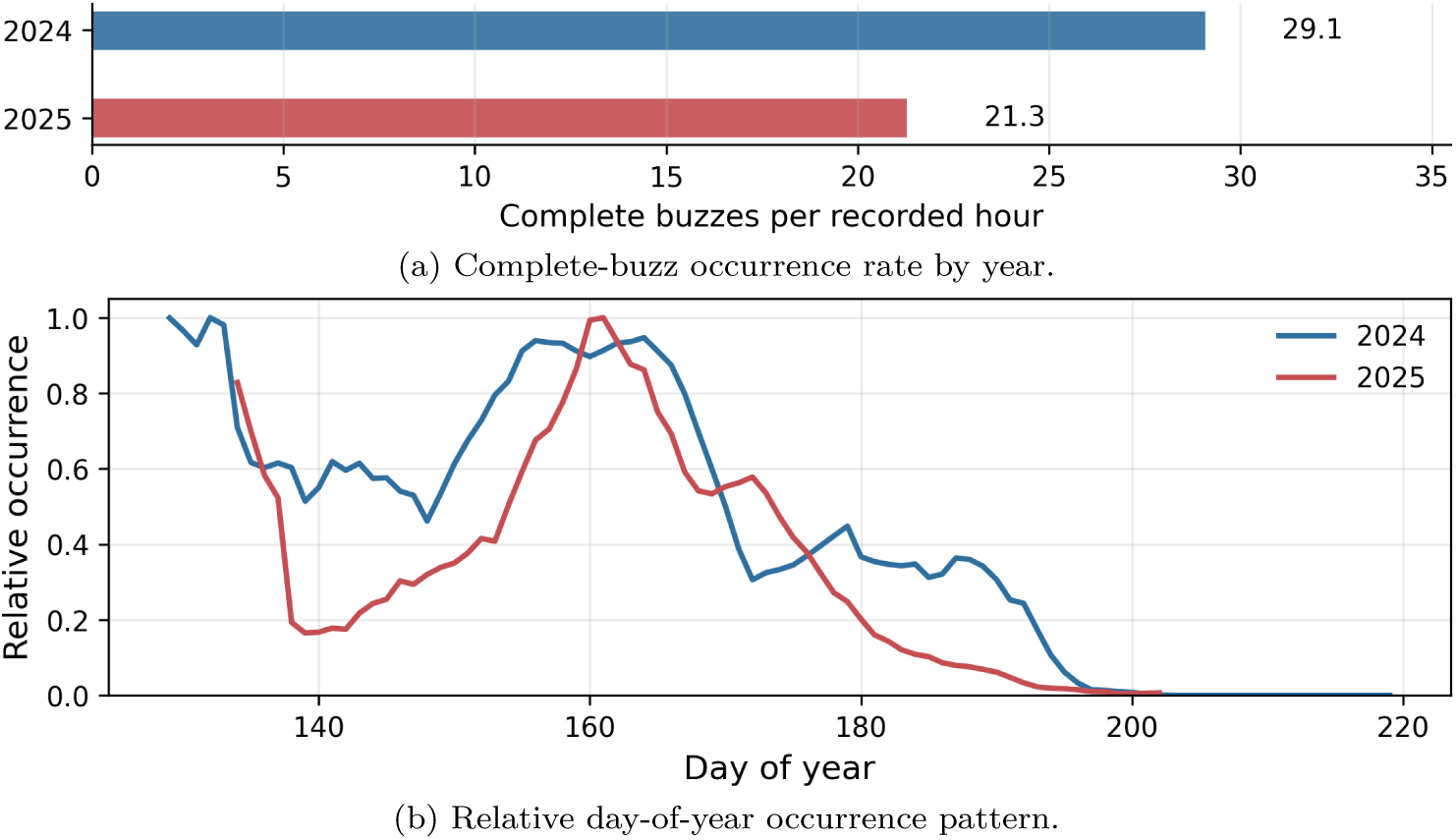
Between-year occurrence patterns from retained complete-buzz records. Panel (a) shows complete buzzes per recorded hour by year. Panel (b) shows relative 7-day smoothed complete-buzz occurrence after scaling each year to its own maximum.

## 9. Discussion

Ecological questions about individual vocalizations require more than fixed-window presence predictions. This study shows that event localization, duplicate-window consolidation, and event-quality curation can turn noisy field recordings into auditable ecological measurements. This structure makes retrieval, boundary placement, duplicate-window handling, and completeness curation separately auditable rather than collapsing them into a single performance score.

The AST and YOLOv11-cls baselines provide a useful window-level benchmark: the SpectroBird detector is modestly lower under complete-within-window scoring, but under the partial-overlap criterion it exceeds AST and is effectively tied with the YOLOv11-cls baseline for complete buzzes. The more important distinction for event-level ecological measurement is output structure. A full-window classifier could serve as an initial screen, but an analysis requiring duration or frequency bounds would still need a second localization step to convert positive windows into event-level time-frequency records, introducing additional annotation, calibration, and error-propagation issues. The detection-localization formulation keeps that localization within the core model while remaining competitive for detection.

Event consolidation is the bridge between tiled spectrogram inference and ecological records. Standard box-merging logic is not sufficient in this setting because overlap has two meanings: repeated detections of the same vocalization in adjacent windows, and genuinely distinct vocalizations that overlap in time and frequency. The consolidation procedure is therefore a domain-specific adaptation of windowed object-detection post-processing. The one-to-one event-target matching protocol serves a similarly specific role: it defines event identity for retrieval evaluation, while localization quality is reported afterward through IoU and boundary-error summaries. Together, these choices prevent plausible same-event detections from being discarded solely because of boundary disagreement, while helping preserve overlapping vocalizations as separate event hypotheses.

The field summaries supported one of the two elevation predictions: complete buzzes were longer at lower elevations, whereas complete-buzz occurrence showed no simple elevation trend. The occurrence result indicates that elevation alone does not explain complete-buzz rates. The decline in duration supports the prediction that males at lower-elevation sites produce longer buzz displays, although the mechanism remains unresolved. Complete-buzz occurrence was descriptively lower in 2025, but the adjusted year effect was not statistically significant. The strongly correlated day-of-year occurrence curves show that the broad seasonal pattern was similar between years.

The winter preceding the 2025 field season was colder and snowier than the preceding winter, with regional reports also documenting cold waves and deficient pre-monsoon rainfall in Chamba (Open-Meteo, 2026; India Meteorological Department, Meteorological Centre Shimla, 2025b,a). These differences provide possible ecological context for the lower aggregate occurrence observed in 2025.

The occurrence, duration, and seasonal results also clarify why event-level outputs matter. The duration result depends on reliable start and end estimates and on filtering incomplete events before aggregation. The occurrence and seasonal summaries depend on event consolidation, because close or overlapping buzzes should contribute separately to rate and timing rather than being reduced to a single positive window. This is why SpectroBird was designed around individual vocal events rather than fixed window-level presence alone.

### 9.1. Limitations and Reproducibility

This study relies on a sequence of fixed processing choices and thresholded post-processing steps. While this enabled rapid methodological iteration, it increases reproducibility burden and motivates future consolidation into a standardized execution protocol. The consolidation procedure was fixed before held-out reporting, but was not exhaustively compared against alternative post-processing methods. Additional limitations include empirical evaluation on a single species and call type, annotation effort requirements, dependence on post-processing thresholds and consolidation parameters, observed error modes such as faint overlaps or structured noise that mimics buzz texture, and annotation uncertainty near event boundaries. Completeness-curation metrics are validation-set estimates rather than results from a separate held-out completeness benchmark. Baseline comparisons are harmonized with a shared window-level scoring protocol, but direct event-localization outputs still involve formulation-specific post-processing for downstream ecological analyses. We therefore document fixed parameters, identify sensitivity checks needed for downstream ecological inference, and retain appendix documentation of adjudication and parameter settings.

## 10. Conclusion

This study was driven by an ecological measurement problem: testing whether HLW buzz occurrence and structure vary across elevation and year required records of individual buzzes. The curated event records produced by SpectroBird supported the motivating ecological questions: complete buzzes were longer at lower elevations, complete-buzz occurrence showed no simple elevation trend, and relative day-of-year occurrence patterns were strongly correlated between years.

The event-record requirement matters because window-level classifiers can estimate broad vocal presence, but they do not preserve event identity, timing, or boundaries. By combining time-frequency localization, overlapping-window consolidation, and completeness filtering, SpectroBird provides a practical template for avian acoustic studies that require curated records of individual vocalizations rather than window-level presence alone.

## Acknowledgments

This research was supported in part through research cyberinfrastructure resources and services provided by the Partnership for an Advanced Computing Environment (PACE) at the Georgia Institute of Technology, Atlanta, Georgia, USA. RRID:SCR_027619.

BGF was supported by a Packard Fellowship for Science and Engineering from the David and Lucile Packard Foundation (2024-77385).

## Data Availability

The model training and test datasets used in this study will be made available upon publication.

## Appendix A. Coordinate Mapping Details

For each analysis-window index *n* and predicted box (*x_i_, y_i_, w_i_, h_i_, c_i_*), the detection-localization model maps normalized detector-image coordinates to absolute time and frequency bounds by

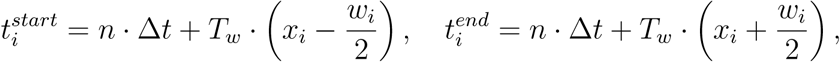

and

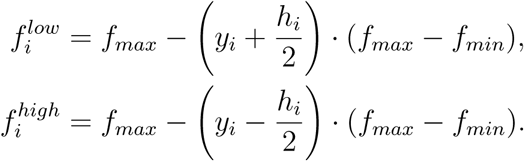

Here Δ*t* is window stride and *T_w_* is window duration. In this study,

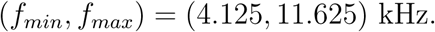

## Appendix B. Event-Consolidation Details

This appendix gives the fixed scoring details for the event-consolidation procedure summarized in Section 5.5. Consolidation is performed in absolute time-frequency coordinates. Each raw detection is represented as

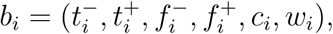

where *t^−^_i_*, *t^+^_i_* are absolute start and end times, *f^−^_i_*, *f^+^_i_* are lower and upper frequency bounds, *c_i_ ∈* [0, 1] is detector confidence, and *w_i_* is the source analysis-window index. Raw detections are treated as nodes in a within-recording association graph. Candidate same-event links are considered only between detections from different but nearby analysis windows within the same recording; detections from the same analysis window are never linked directly for consolidation:

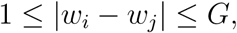

with fixed *G* given in Table C.1.

The output of consolidation is a set of event tracks,

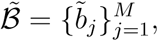

where each *b^~^_j_* represents one candidate buzz event in absolute time-frequency coordinates.

Figure B.1 shows the duplicate-window case for a single buzz before and after consolidation.

**Figure B.1:**
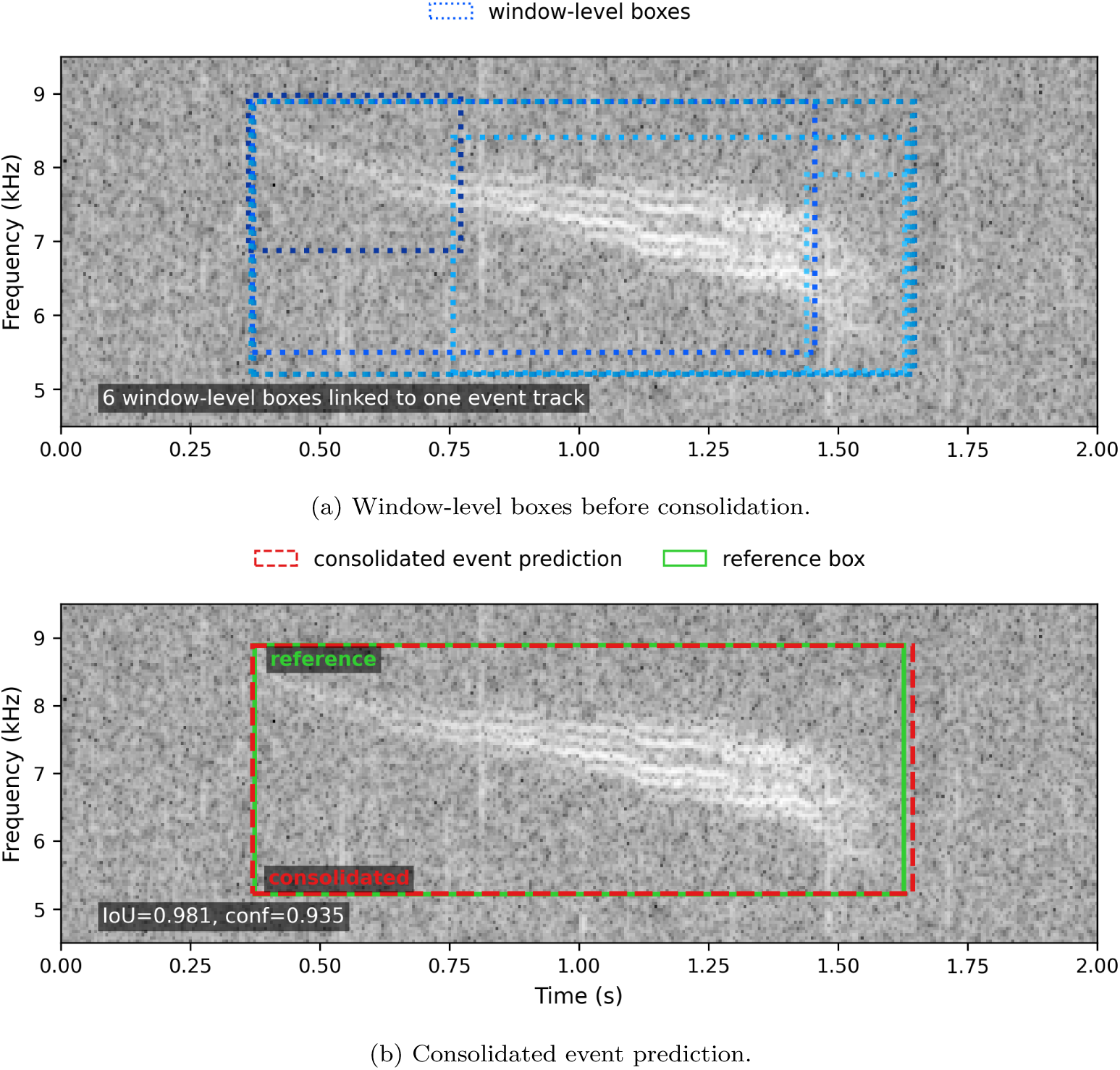
Windowed prediction consolidation for a single buzz. Window-level boxes from adjacent overlapping windows are consolidated into one event-level prediction, shown with the reference box for comparison.

For a candidate link (*b_i_, b_j_*), let

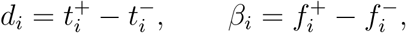

and let *m_t_*(*i*) and *m_f_* (*i*) denote the time and frequency centers of *b_i_*. Duration and bandwidth similarity are

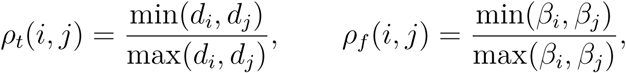

and normalized center offsets are

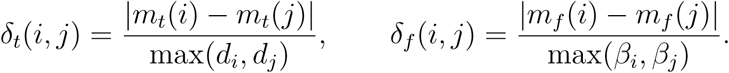

To quantify whether a detection is likely truncated by its source window, let *ℓ_i_* and *r_i_* denote the normalized left and right time coordinates of *b_i_* within that window. Its edge proximity is

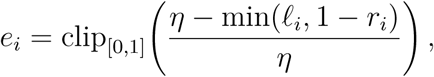

where *η* is the fixed edge-proximity scale. For a candidate link, we use the larger edge-proximity value,

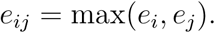

The pairwise association score is

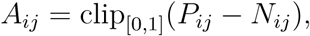

The fixed weights *a*_1_*, …, a*_10_ are reported in Table C.1. Here IoU_2*D*_ is the full time-frequency box IoU, while IoU*_t_* and IoU*_f_* are one-dimensional IoU on the time and frequency intervals.

A candidate link is retained only if it is *mutual-best* under *A_ij_*, meaning that *b_j_* is the highest-affinity partner of *b_i_* and *b_i_* is the highest-affinity partner of *b_j_*. Let

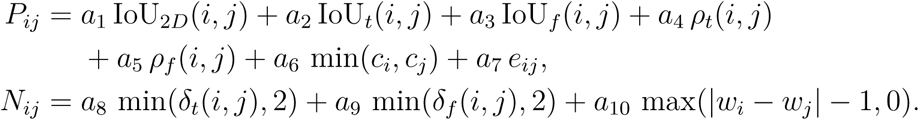

denote the mutual-best margin over the second-best alternative on each side. A mutual-best pair is retained only if it passes one of two fixed gates, a *strong-link* gate or a *support-link* gate (Table C.1). Strong links seed event tracks. Support links are processed afterward and may extend a seeded track, but they are not allowed to merge two already established tracks. To preserve distinct overlapping buzzes as separate event hypotheses, an event track may contain at most one detection from any given analysis window.

The procedure also treats window-edge fragments explicitly. After the initial linking pass, a near-edge singleton detection *p* may be absorbed into an existing multi-window track *T_k_* with absolute time-frequency envelope

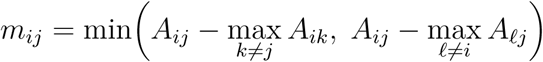

Containment of *p* within *E_k_* is summarized by area, temporal, and frequency cover quantities. Area cover is *|p ∩ E_k_|/|p|*, time cover is the fraction of the singleton’s temporal extent contained in *E_k_*, and frequency cover is the fraction of the singleton’s frequency extent contained in *E_k_*:

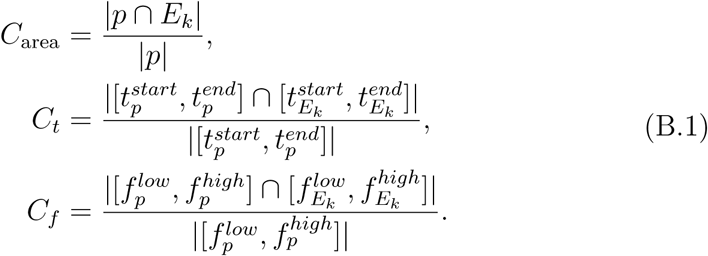

Using those terms, the singleton-to-track absorption score is

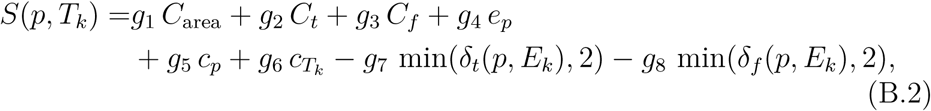

where *c_T_* = max*_i_*_∈*T*_ *c_i_* is the maximum member confidence of the candidate target track, and *δ_t_*(*p, E_k_*) and *δ_f_* (*p, E_k_*) are normalized center offsets computed analogously to the pairwise definitions above after replacing the second box with the track envelope. The fixed weights *g*_1_*, …, g*_8_ are reported in Table C.1. A singleton is absorbed only if it is sufficiently edge-adjacent, the target track does not already contain a detection from the singleton’s source window, and the fixed containment, score, and assignment-margin gates are all satisfied.

Finally, each accepted track *T_k_* is converted to a consolidated event prediction ^~^*b_k_*. Frequency bounds are estimated by confidence-weighted medians of member-detection boundaries,

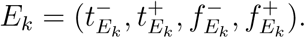

Time bounds use the same weighted-median idea, but with edge censoring: starts near the left edge of a source window are excluded from the start vote, and ends near the right edge are excluded from the end vote, so that potentially truncated partial views do not pull the consolidated boundary inward. These censored observations can still preserve observed extent by expanding the final boundary when necessary. The consolidated event confidence is

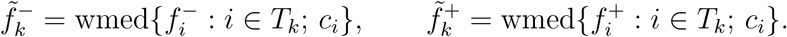

No event-confidence threshold is applied during consolidation itself; low-confidence singleton events remain available for downstream evaluation and export thresholding.

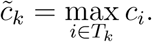

## Appendix C. Parameter Tables

Tables C.1–C.3 summarize the fixed event-consolidation, event-target matching, and completeness-curation configuration settings used in this study.

**Table C.1:** Event-consolidation parameter summary (fixed configuration used in this study).

| Parameter | Value |
| --- | --- |
| Raw prediction ingestion | Window-level detector boxes with confidence $c \geq 0.001$ |
| Candidate window gap | Up to 3 analysis-window steps |
| Association features | 2D, time, and frequency IoU; center distances; duration/bandwidth similarity; minimum confidence; time-edge proximity; window gap |
| Association score weights | 0.45 2D IoU; 0.14 time IoU; 0.14 frequency IoU; 0.10 duration ratio; 0.08 bandwidth ratio; 0.05 minimum confidence; 0.06 edge support; subtracting 0.05 center-time distance, 0.04 center-frequency distance, and 0.03 per extra window gap |
| Strong-link gate | affinity $\geq 0.72$, 2D IoU $\geq 0.55$, minimum confidence $\geq 0.25$, mutual-best margin $\geq 0.04$ |
| Support-link gate | affinity $\geq 0.62$, 2D IoU $\geq 0.35$, minimum confidence $\geq 0.10$, mutual-best margin $\geq 0.03$, window gap $\leq 2$ |
| Support-link requirements | Candidate edge: maximum confidence $\geq 0.70$ or edge support $\geq 0.50$; track merge: one side has maximum confidence $\geq 0.70$ or an existing strong edge |
| Edge-proximity scale | 0.08 of analysis-window duration |
| Edge-singleton absorption | enabled for near-edge singleton detections mostly contained in established tracks |
| Absorption edge requirement | Edge proximity $\geq 0.80$ for singleton detections |
| Absorption containment thresholds | area cover $\geq 0.70$, time cover $\geq 0.55$, frequency cover $\geq 0.55$ |
| Absorption score threshold | score $\geq 0.72$ with margin $\geq 0.08$ |
| Absorption score weights | 0.55 area cover; 0.15 time cover; 0.15 frequency cover; 0.08 edge value; 0.04 singleton confidence; 0.03 target-track confidence; subtracting 0.04 center-time distance and 0.03 center-frequency distance |
| Overlap-preservation constraint | At most one detection from a given analysis window per event track |
| Event-boundary fusion | Time: edge-censored confidence-weighted median with extent preservation; frequency: confidence-weighted median |
| Event confidence | Maximum member-detection confidence |

**Table C.2:** Event-target matching protocol for direct event-level evaluation on the final held-out target labels.

| Parameter | Value |
| --- | --- |
| Matching scope | Within recording |
| Development data | Detection-localization training/validation predictions and labels; fixed before final held-out reporting |
| Prediction unit | Consolidated event prediction |
| Reference unit | Final adjudicated time-frequency target box |
| Eligible predictions | All consolidated predictions are available to the matching sweep before applying the current event-confidence threshold |
| Candidate gate | Time overlap or center-time distance $\leq 0.50$ s; frequency overlap or center-frequency distance $\leq 1.0$ kHz |
| Assignment method | One-to-one linear-sum assignment (Hungarian matching) |
| Assignment score | $\text{IoU}_{2D} + 0.25\text{IoU}_t + 0.25\text{IoU}_f + 0.25C_{\text{small}} - 0.10\delta_t - 0.10\delta_f + 0.001c$, where $C_{\text{small}}$ is smaller-box coverage, $\delta_t$ and $\delta_f$ are normalized center-distance penalties, and $c$ is event confidence |
| IoU interpretation | Localization metric reported after matching; not a hard event-identity gate |
| True positives | Assigned target-prediction pairs |
| False negatives | Unassigned reference targets |
| False positives | Eligible unassigned consolidated predictions; for stratum-restricted rows, candidate matches to the opposite-stratum ignore set are excluded |
| PR-threshold handling | Filter predictions at each confidence threshold, then rerun assignment |
| Calibration diagnostic | All 716 validation labels had a matched prediction at $\text{IoU}_{2D} \geq 0.50$ when predictions were saved at confidence $\geq 0.001$ |
| Low-IoU handling | Plausible same-event matches below $\text{IoU}_{2D} = 0.50$ are retained and evaluated through localization-error summaries |

**Table C.3:** Completeness-curation parameter summary (configuration used in this study).

| Parameter | Value |
| --- | --- |
| Classifier input size | $288 \times 288$ |
| Crop construction | Reconstructed from source audio using consolidated event bounds; predicted event right-aligned in the standardized temporal crop |
| Temporal target crop length | 288 frames |
| Image standardization | Aspect-preserving resize followed by padding to $288 \times 288$ |
| Padding strategy | Local temporal background during labeling; mean/gray padding for square image standardization |
| Classifier model family | YOLOv11-cls |
| Checkpoint selection | Highest validation peak F1 on $D_{\text{cur-val}}$ |
| Completeness score | $p(\text{complete})$ |
| Completeness threshold | $\theta_{\text{cur}} = 0.530$ from validation peak F1 |
| Export eligibility | Applied after the validation-derived detection event-confidence operating threshold |
| Training epochs | 100 |
| Training batch size | 32 |
| Output classes | complete, incomplete |

## Appendix D. Annotation and Agreement Details

This appendix records annotation quality-control procedures for transparency and reproducibility. These procedures are supporting procedural details and are not presented as a novel contribution of this paper.

Procedure summary

- Two independent researcher passes were completed on the held-out test recordings.
- An expert adjudication pass reviewed candidate buzz events and boundary placement for the held-out event-detection test set.
- Agreement was quantified over timeline bins including silence intervals.

### Adjudication rule (supporting detail)

The annotation protocol used weighted consolidation: expert annotations carry higher decision weight than individual non-expert annotations. Events supported by the expert and at least one researcher are accepted; events unsupported by the expert are not promoted solely by non-expert agreement. We include this here as provenance of label curation logic rather than as a methodological claim.

### Agreement statistic

Krippendorff’s alpha for the finalized test annotation timeline was 0.92, computed over timeline bins including silence intervals.

## Appendix E. Diagnostic Error Cases

**Figure E.1:**
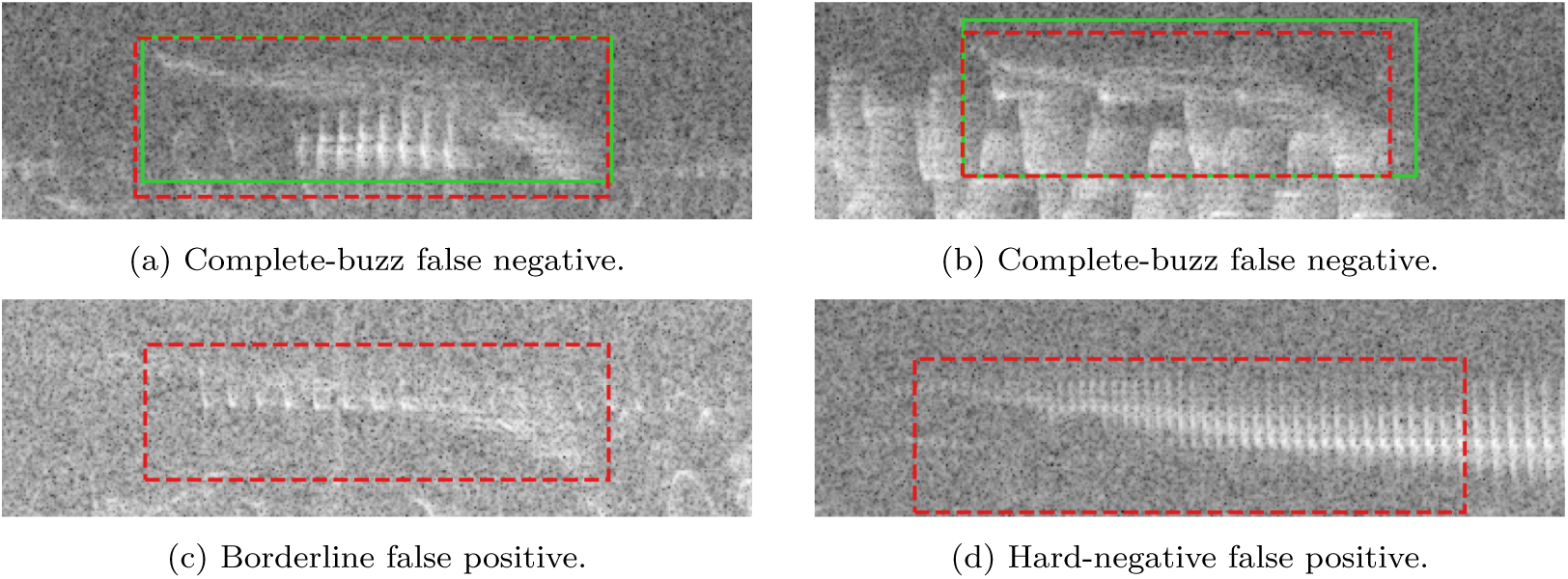
Representative detector error cases at the held-out peak-F1 reporting threshold for complete buzzes (*θ* = 0.838761). Panels (a) and (b) show false negatives on complete buzzes in which the model still produced well-localized consolidated event predictions (dashed red) below this reporting threshold; the consolidated event confidences were 0.791 and 0.798, respectively, and reference boxes are shown in solid green. Panel (c) shows a borderline buzz-like case in which non-stationary interference overlaps a faint buzz-like trace, yielding a false positive consolidated event. Panel (d) shows a hard negative case with no annotated buzz event, but with high-band structure resembling a buzz and therefore producing a false positive prediction. In all panels, time increases from left to right and frequency from bottom to top.

## Appendix F. Localization Distributions

**Figure F.1:**
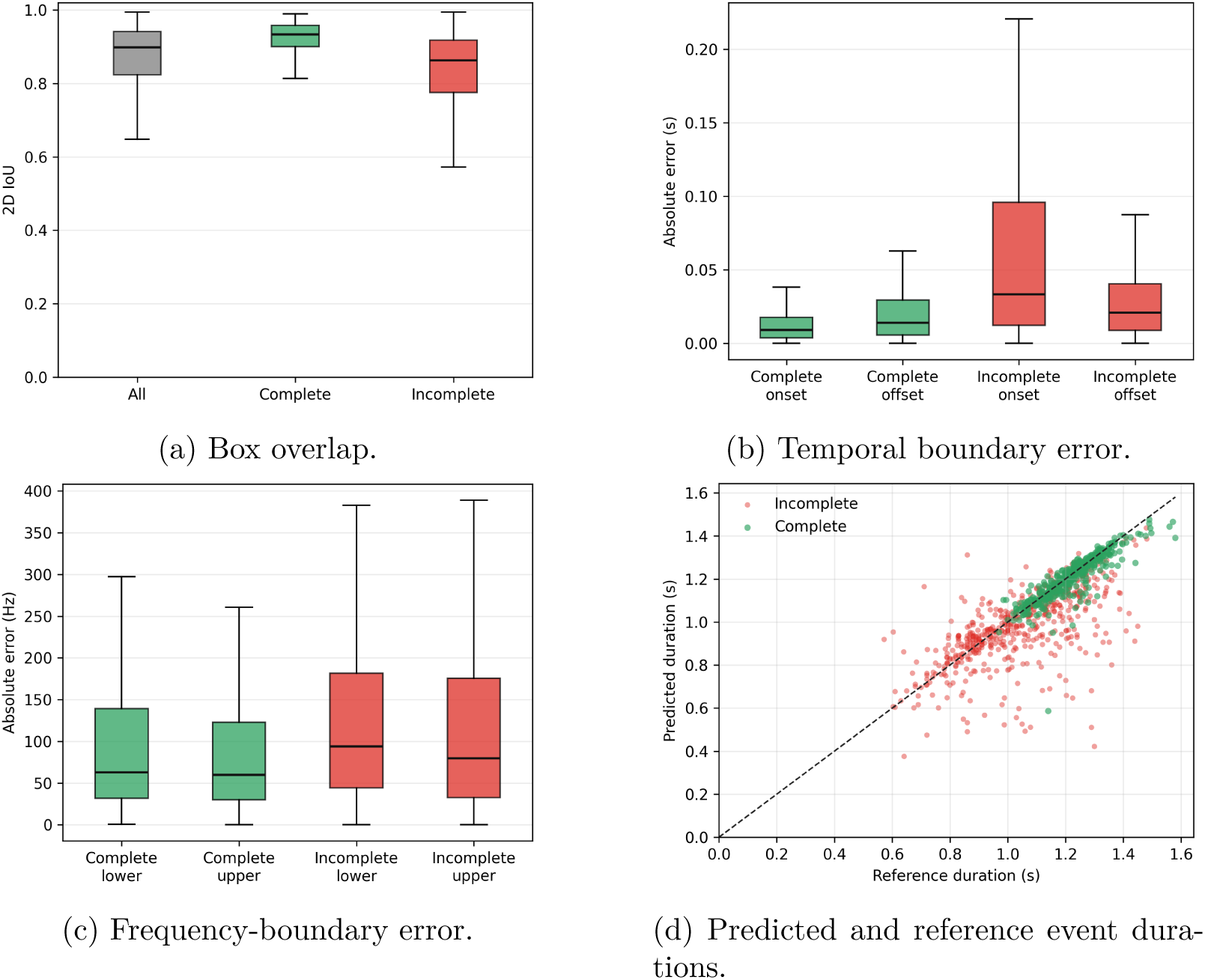
Detection-localization distributions for matched true-positive consolidated events at the all-buzz peak-F1 event-confidence threshold.

